# De novo multi-mechanism antimicrobial peptide design via multimodal deep learning

**DOI:** 10.1101/2024.01.02.573846

**Authors:** Yue Wang, Haifan Gong, Xiaojuan Li, Lixiang Li, Yinuo Zhao, Peijing Bao, Qingzhou Kong, Boyao Wan, Yumeng Zhang, Jinghui Zhang, Jiekun Ni, Zhongxue Han, Xueping Nan, Kunping Ju, Longfei Sun, Huijun Chang, Mengqi Zheng, Yanbo Yu, Xiaoyun Yang, Xiuli Zuo, Yanqing Li

## Abstract

Artificial intelligence (AI)-driven discovery of antimicrobial peptides (AMPs) is yet to fully utilise their three-dimensional (3D) structural characteristics, microbial specie-specific antimicrobial activities and mechanisms. Here, we constructed a QLAPD database comprising the sequence, structures and antimicrobial properties of 12,914 AMPs. QLAPD underlies a multimodal, multitask, multilabel, and conditionally controlled AMP discovery (M3-CAD) pipeline, which is proposed for the *de novo* design of multi-mechanism AMPs to combat multidrug-resistant organisms (MDROs). This pipeline integrates the generation, regression, and classification modules, using a innovative 3D voxel coloring method to capture the nuanced physicochemical context of amino acids, significantly enhancing structural characterizations. QL-AMP-1, discovered by M3-CAD, which possesses four antimicrobial mechanisms, exhibited low toxicity and significant activity against MDROs. The skin wound infection model demonstrates its considerable antimicrobial effects and negligible toxicity. Altogether, integrating 3D features, specie-specific antimicrobial activities and mechanisms enhanced AI-driven AMP discovery, making the M3-CAD pipeline a viable tool for *de novo* AMP design.

## MAIN

The escalating threat posed by multidrug-resistant organisms (MDROs) to human health necessitates an urgent search for new antimicrobial solutions^1, 2^. As antimicrobial resistance (AMR) pervades all clinically used antibiotics, developing substitutes with multiple novel mechanisms is crucial^3–6^.

Antimicrobial peptides (AMPs) are key players in innate immunity and provide primary defence against pathogens^7, 8^. Their amphiphilic and positively charged nature allows for the selective disruption of negative bacterial membranes while sparing the neutral mammalian cell membranes^9, 10^. This nonspecific antibacterial action based on the biophysical interactions reduces the risk of inducing AMR^11–13^, which positions AMPs as potential next-generation antimicrobials and as promising alternatives to traditional antibiotics^14^. However, traditional AMP discovery using wet experiments is time-consuming and costly^15, 16^. Consequently, artificial intelligence (AI)-driven approaches such as identification^17–21^, optimization^22–24^, and generation^16, 25–28^ have been proposed to accelerate AMP discovery. Among these, generative models overcome the limitations of identification and optimisation models. Unlike identification models, which screen targets within a defined peptide range, and optimisation models, which require manual selection of template sequences, generative models theoretically yield optimal AMPs that meet the specific target conditions.

All these AI-driven AMP discovery models require large-scale, manually labelled data for empowerment. Driven by the significance of training data, the following question arises: What human knowledge and peptide features can be used to enhance the inhibitory activity of the AMPs discovered by AI, particularly generative AI, against superbugs?

Our literature review revealed the following findings. First, the current AI-driven AMP discovery predominantly utilises limited human knowledge, primarily regarding whether a peptide exhibits antibacterial properties, often neglecting the varying efficacy of AMPs against distinct bacteria, particularly clinically isolated MDROs^29^. Second, most existing studies assume that the AMPs function by binding to and disrupting negatively charged bacterial membranes, thereby adopting membrane disruption-related features in the AMP discovery frameworks^17, 21^ and verifying the AMPs’ membrane permeabilisation and depolarisation abilities through experimental methods^16–21, 24, 27, 28^. This resulted in other antimicrobial mechanisms such as the inhibition of biofilm formation, DNA replication, and protein synthesis to be overlooked during the design phase. Finally, the existing studies rely heavily on primary sequence data, neglecting crucial three-dimensional (3D) structural information that reflects secondary and tertiary structures, which have major significance in peptide function^29^. However, recent advancements in AI models such as AlphaFold2 and RoseTTAFold have improved the precision of peptide and protein structure prediction^30, 31^, paving the way for enhanced peptide characterisation and downstream tasks using 3D structural data.

We addressed these gaps and curated a comprehensive AMP database (QLAPD) by meticulously selecting 12,914 AMP entries from existing knowledge bases^32–35^ and numerous published papers. Each entry recorded eight feature categories, including the predicted 3D structure, microbial specie-specific antimicrobial activities, antimicrobial mechanisms, and toxicity. We propose a multimodal, multitask, multilabel, and conditionally controlled AMP discovery (M3-CAD) pipeline to generate and optimise new AMPs with multiple antimicrobial mechanisms, broad-spectrum resistance to MDROs, and low host toxicity. We also introduce a 3D voxel coloring method and a multilabel rebalancing training strategy to address the challenges in representing 3D peptide structures and the long-tail distribution problem in multilabel classification tasks.

Wet lab validation of the M3-CAD selected candidate sequences demonstrated their inhibitory abilities against 26 pathogenic bacterial strains. The most effective peptide, QL-AMP-1, displayed powerful antimicrobial activity against gram-positive and gram-negative MDROs, low host toxicity, and four antimicrobial mechanisms. Unlike clindamycin, QL-AMP-1 did not induce significant resistance after 20 passages of bacterial cultivation. QL-AMP-1 significantly reduced the bacterial load with no observable *in vivo* toxicity in full-thickness skin wound models that were infected with *S. aureus* and *A. baumannii*. Comprehensively, incorporating the peptide 3D structural features and information on the antimicrobial mechanism enhanced AI-driven AMP discovery, rendering the M3-CAD pipeline a promising tool for *de novo* AMP design.

## RESULTS

### QLAPD: A database that links the sequences, structures, and properties of AMPs

QLAPD, an AMP database, reviewed and annotated by clinical doctors and microbiology experts from Qilu Hospital, was established to train an AI framework capable of *de novo* design of AMPs that have multiple antimicrobial mechanisms and inhibit MDROs. QLAPD encompasses 12,914 AMPs and each records its amino acid sequence, predicted 3D structure, and six functional attributes, namely: (1) inhibitory activity against six drug-resistant pathogenic bacteria; (2) the number of drug-resistant bacteria it can inhibit; (3) the number of non-resistant bacteria it can inhibit; (4) four antimicrobial mechanisms such as disrupting bacterial membranes; (5) toxicity; and (6) nephrotoxicity among the six kinds of organ toxicity (Fig. 1a, Supplementary Fig. 1a-f). The 3D structure of the peptide was predicted using the AlphaFold2 model^30^ and stored in the Protein Data Bank (PDB)^36^, which recorded the type of each atom in the peptide, coordinates of the atom in 3D space, and the type of amino acid residue to which the atom belongs (Fig. 1a).

**Fig. 1.**
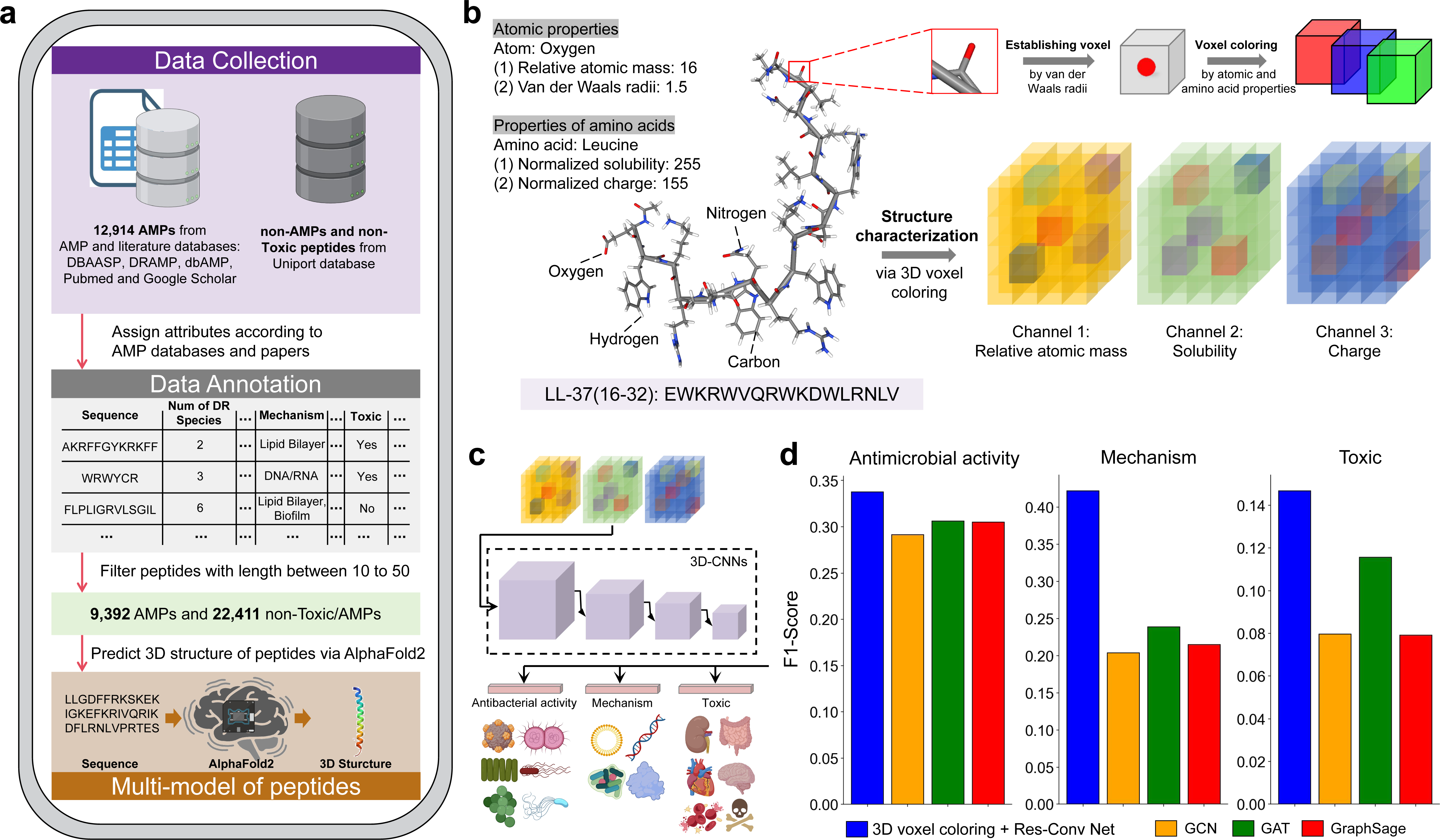
Overview of the proposed QLAPD database and 3D voxel coloring method. **a,** Schematic flow chart of antimicrobial peptide collection, data cleaning, label annotation and multi-modal feature construction of QLAPD database. **b,** Schematic diagram of the 3D voxel coloring method, taking the antimicrobial peptide LL-37(16-32) as an example. **c,** The schematic diagram shows that the 3D structural features of polypeptides obtained using the 3D voxel coloring method are used to train three downstream multi-label classification tasks of the 3D-CNN model. **d,** The bar plots shows the performance of the 3D voxel coloring + Res-Conv Net and three graph neural network methods in the multi-label classification task of antimicrobial activity, mechanism and toxicity of antimicrobial peptides. Shown are the means obtained by five-fold cross validation.

### 3D Voxel coloring to improve structural characterisation of the peptides

Inspired by the solutions to 3D visual tasks, a 3D voxel coloring method was proposed to improve the 3D structural representation of the peptides. A peptide was placed in a 3D Cartesian coordinate system with its centroid as the origin, and housed within a cube comprising equal 3D voxels. The occupied voxels were determined by the spatial location of atoms (predicted by AlphaFold2^30^) and their van der Waals radii^37^. The voxels within the atom-covered area have feature channel-assigned values based on the properties of the corresponding atoms, defined here by the atomic mass, solubility (indicating hydrophobicity), and acidity or alkalinity (indicating charge). The unoccupied 3D voxels were assigned a value of zero (Fig. 1b, Supplementary Table 1). This paradigm allows us to adeptly convert the complex challenge of peptide 3D structural characterization into a tractable 3D visual feature extraction task, and solved by introducing 3D convolutional neural network (3D-CNN) models (Fig. 1c).

To assess this method, we compared the 3D-CNN performance, based on the 3D voxel coloring method (3D Res-Conv Net^38^, 3D SwinUNETR Net^39^), against traditionally used Graph Neural Networks (GNNs)^40^, such as GCN^41^, GAT^42^, and GraphSage^43^, for the multilabel classification of peptide antimicrobial activity, mechanisms, and toxicity. The five-fold cross-validation results suggest that the 3D-CNN models outperformed the GNNs in most metrics across the three downstream tasks, with 3D Res-Conv Net having the highest overall score. (Fig. 1d and Supplementary Tables 2-4).

### M3-CAD pipeline design

We proposed an M3-CAD pipeline that consists of sequential generation, regression, and classification modules (Fig. 2a-c). Compared to previous studies, the greatest advantage of M3-CAD is that it uses the antimicrobial mechanism data and inhibitory activity data of the AMPs against MDROs during training, thereby learning the potential relationship between the sequence and structure of peptides and the functional attributes, such as four antimicrobial mechanisms and inhibitory ability against six types of drug-resistant bacteria.

**Fig. 2.**
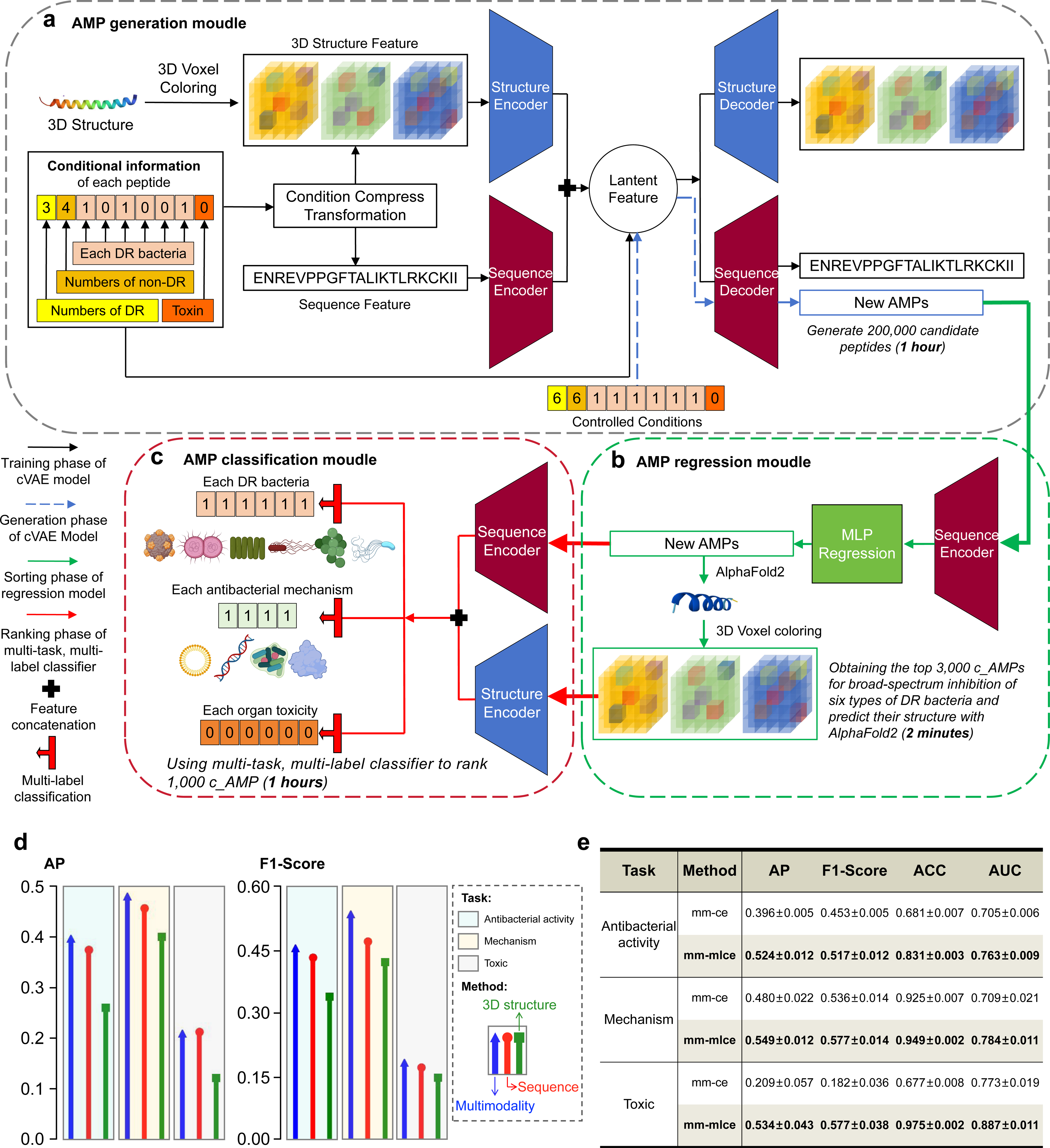
Workflow and performance evaluation of the M3-CAD pipeline. **a-c,** Schematic showing the generation (a), regression (b) and classification (c) moudles that make up the M3-CAD pipeline. **d,** The lollipop plot demonstrates the performance of multi-modality-based models and models based only on sequence or 3D structural modalities on the multi-label classification task of antimicrobial activity, mechanism and toxicity. Shown are the means obtained by five-fold cross validation. **e,** The table shows the performance improvement of three downstream tasks of AMPs by applying the multi-label rebalancing loss function (mm-mlce). Shown are the means and standard deviations obtained by five-fold cross validation.

The training and cross-validation data for M3-CAD included 9,392 AMPs with lengths of 10-50 amino acid residues selected from the QLAPD database, and 22,411 non-AMPs of the same length range collected from the Uniport database^44^. The six functional attributes of these peptides were defined as multilabel one-hot encoding (activity against six drug-resistant bacteria, toxicity to six types of organs) or multiclass labels (number of drug-resistant bacteria inhibited, number of non-drug-resistant bacteria inhibited, and toxicity) for generation, regression, and classification tasks (Figs. 2a and c). In practice, generating 200,000 new candidate AMPs (c_AMPs) and further screening of high-priority AMPs, using the trained M3-CAD pipeline, required approximately 2 h, greatly accelerating the discovery of AMPs.

### Generating AMPs that satisfy the given properties based on a multimodal VAE

The centerpiece of M3-CAD is a generative model based on a multi-modal, condition-controlled variational autoencoder^45^ (cVAE) designed to generate new peptides that meet the specified attributes (Fig. 2a). Unlike previous VAEs, this cVAE incorporates two encoders and decoders for the sequence and 3D structure modalities. For sequence features, peptides up to 50 residues long were encoded into a 1×50 dimensional tensor and feature extraction was performed using a multi-layer perceptron (MLP). The 3D structural features employed the 3D voxel coloring method and 3D Res-Conv Net. To integrate sequence and structural information with functional attribute information, the four functional attributes of the peptide were represented as a 1×9 dimensional tensor, feature-extracted using MLP, and compressed to a 1×1 tensor. This tensor was multiplied by the sequence and structural features to input into the encoder, and concatenated with the latent feature between the encoder and decoder, allowing the cVAE to learn the relationship between peptide sequences, structural features, and controlled conditions. During the inference phase, the trained decoder sampled peptides that met the conditions from the latent-space probability distribution by inputting the corresponding conditions (Fig. 2a).

In practice, using the proposed cVAE, approximately 200,000 novel c_AMPs can be designed within 1 h, which potentially satisfy various antimicrobial mechanisms and broad-spectrum inhibitory conditions against MDROs, thereby greatly expanding the combinatorial molecular space for AMP searches.

### Ranking the generated peptides using an MLP regression model

Given the time-intensive nature of predicting the 3D structures of numerous peptides, we incorporated a sequence-based MLP as the second component of M3-CAD for the initial screening of the generated sequences (Fig. 2b). The MLP is trained to rank peptides based on their ability to broadly inhibit drug-resistant bacteria, and a higher priority is given to broad-spectrum inhibitory peptides. In practice, this stage of M3-CAD selects 3,000 high-priority c_AMPs for 3D prediction using AlphaFold2, progressing them to the final stage of the pipeline. A comparison between the predicted antimicrobial activity of the 3,000 MLP-selected c_AMPs and 3,000 randomly sampled peptides from cVAE confirmed the superior performance of the MLP-selected c_AMPs in a broader spectrum of antimicrobial activity (Supplementary Fig. 2a).

### Identifying the attributes of AMPs using a multilabel classification model trained by the rebalancing loss function

The final component of M3-CAD is a multimodal, multitask, and multilabel classifier designed to identify AMPs with diverse antimicrobial mechanisms, broad-spectrum inhibition against drug-resistant bacteria, and low toxicity. This classifier employs an MLP and a 3D Res-Conv Net for feature extraction from the peptide sequences and 3D structures in parallel with the previously mentioned cVAE. The extracted features were fused and dispatched to the three classification heads for multilabel classification (Fig. 2c). We evaluated the performance of the multilabel classifier on antimicrobial activity, mechanism, and toxicity prediction tasks, which demonstrated the superior performance of the multimodal model compared to models using only sequence or 3D structure modalities (Fig. 2d and Supplementary Table 5).

During classifier training, we encountered severe label imbalances across the three multilabel classification tasks (Supplementary Fig. 1b-e), which could hamper the performance of the deep neural network model^46^. This prompted us to propose a multilabel rebalancing loss function, by extending the "Softmax + Cross Entropy" scheme to multilabel classification scenarios with multiple target classes. This method ensured that the score of each target class was not lower than that of the non-target class, thereby providing a balanced loss function to handle the imbalanced data distribution^47^. Ablation studies show that the rebalancing learning strategy improved the performance across all three multilabel classification tasks (Fig. 2e).

In this application, 3,000 c-AMPs were ranked based on the predicted antimicrobial activity and toxicity confidence levels from the multilabel classifier. The 10 top-ranked peptides were advanced as the final output of the M3-CAD pipeline for wet lab experiments.

### Confirming the necessity of each component of M3-CAD using wet lab experiments

To validate the necessity of each individual module in the M3-CAD framework, we systematically evaluated the antimicrobial efficacy of the 10 top-ranked peptides that were predicted under varying conditions. The generative module (G), combined generative and regression modules (G+R), and integrated generative, regression, and classification modules (G+R+C) were solely employed for this purpose. This comparative analysis targeted the experimental minimum inhibitory concentration (MIC) values of the peptides against established strains of *S. aureus*, *E. coli*, and *A. baumannii*, as shown in Fig. 3a-c and Supplemental Table 6. The investigation yielded compelling insights. The generative module on its own, devoid of any ranking capability intrinsic to its design, produced peptides that lacked significant antimicrobial properties and had MIC values exceeding 256 µg/ml across a random sample of 10 peptides from a pool of 200,000 generated by the cVAE algorithm. Conversely, incorporating the regression module to prioritise the synthetic candidates resulted in a marked improvement. 70% of the peptides manifested bacteriostatic activity against *S. aureus* and a substantial fraction (60%) exhibited MICs at or below 64 µg/ml. Integrating a multimodal classifier within the M3-CAD paradigm further elevated the performance benchmark and culminated in identifying peptides with consistent inhibitory effects against *S. aureus* in the entire cohort of the 10 peptides that were tested. Remarkably, among these, four peptides showed significant antimicrobial activity with MICs registering at 8 µg/ml, underscoring the enhanced predictive power and potential therapeutic applicability of the M3-CAD system.

**Fig. 3.**
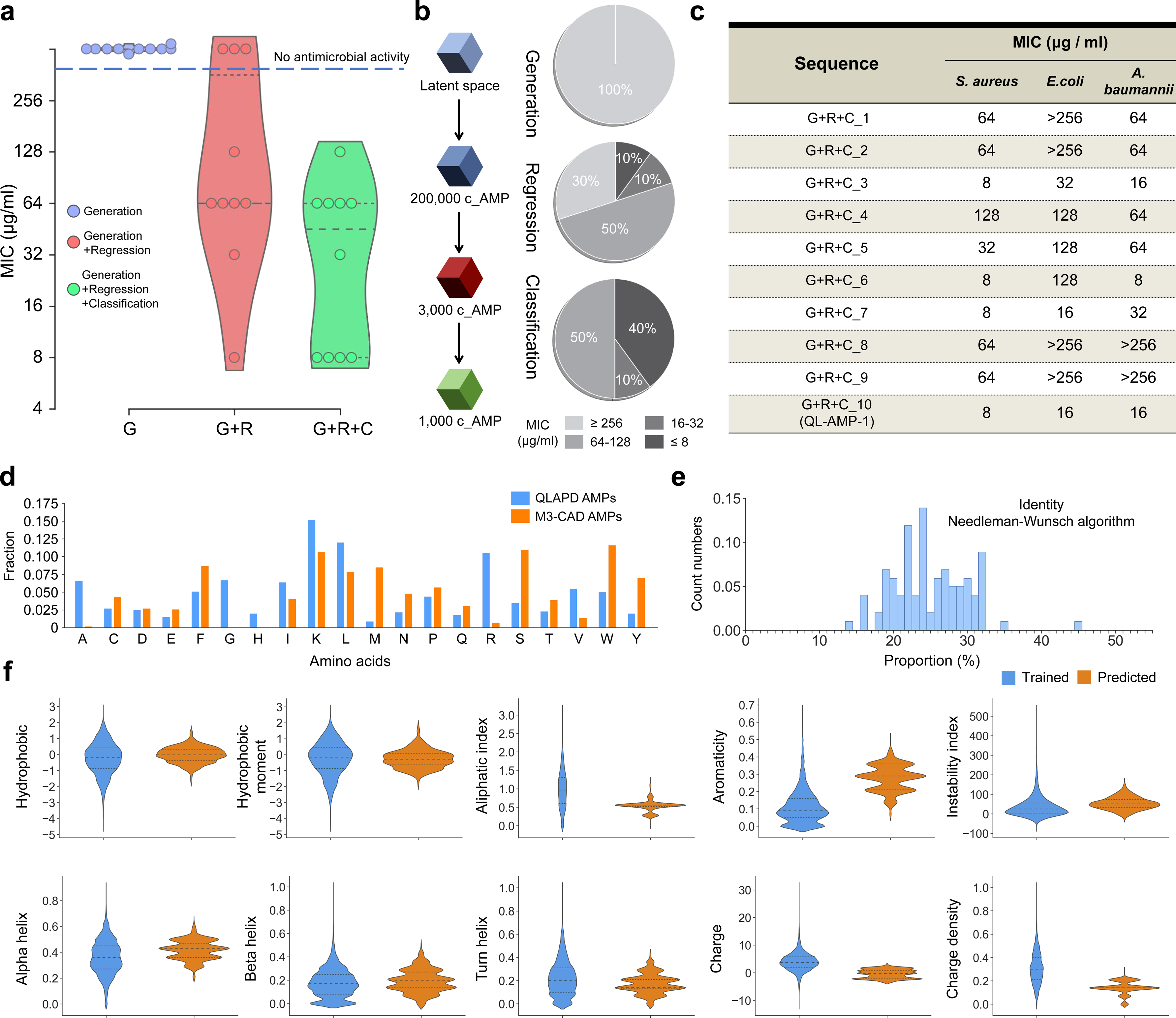
Wet-lab validation of the antimicrobial activity of AMPs discovered by the M3-CAD pipeline and characterization of the physicochemical properties. **a,** Violin plots showing the distribution of the tested MIC values against *S. aureus* CMCC26003 of the top-10 peptides from the complete and partial component pipelines (n = 3 biologically independent replicates). The dashed line represents MIC > 256 µg/mL. The dotted lines in each data group represent the first quartile, median and third quartile. **b,** Overview of the antimicrobial activity distribution against *S. aureus* CMCC26003 of the top-10 peptides from pipelines with different modules. The addition of each module improved the antimicrobial activity of the discovered AMPs (n = 3 biologically independent replicates). **c,** Experimental MIC values against *S. aureus* CMCC26003, *E.coli* CICC21530 and *A. baumannii* ATCC19606 (n = 3 biologically independent replicates) of the top-10 peptides obtained by the complete M3-CAD pipeline. **d,** Comparison of the amino acid compositions of the top-1,000 AMPs discovered by M3-CAD with the AMPs in the QLAPD database (n = 9,392). **e,** Distribution of the highest similarities of the top-1,000 new discovered AMPs by M3-CAD to that of QLAPD. Most of the predicted AMPs have less than 32% similarity to the previously known AMPs. **f,** Comparison of the physicochemical properties of QLAPD-AMPs (n = 9,392) and newly discovered AMPs (n = 1,000) using the M3-CAD pipeline.

### Properties of the AMPs designed by M3-CAD pipeline

We compared the sequence similarity and key molecular features (such as amino acid composition, charge count, hydrophobicity, and hydrophobic moment) of the top 1,000 c_AMPs, produced by the M3-CAD pipeline in a single run, with the AMPs in the training set. Compared to the AMPs in the training set, the peptides designed by M3-CAD exhibit low sequence similarity and display enhanced hydrophobicity and aromaticity, coupled with reduced positive charge, charge density, and a lower aliphatic index. These physicochemical property differences may be attributed to the objective of M3-CAD to design AMPs with multiple mechanisms, as opposed to the AMPs in the training set, which predominantly operate through the membrane disruption mechanism (Fig. 3d-f, Supplementary Table 7).

### *In vitro* evaluation of the leading AMPs designed by M3-CAD

The 10 top-ranked peptides output by the M3-CAD pipeline in a single run were synthesised and tested for their MICs against standard strains of *S. aureus*, *E. coli*, and *A. baumannii*. All peptides inhibited at least one strain at less than 256 μg/mL (Fig. 3c), supporting the AMP design capabilities of M3-CAD. The most efficacious, QL-AMP-1, demonstrated MICs less than or equal to 16 μg/mL against all tested standard strains (Fig. 3c). QL-AMP-1 shared less than 30% sequence similarity with known AMPs, indicating its novelty (Supplementary Table 8). Subsequently, wet lab experiments were performed to explore the antimicrobial activity of QL-AMP-1 against MDROs, off-target toxicity, induced resistance, and antimicrobial mechanism.

### Antimicrobial activities against MDROs

The antimicrobial activity of QL-AMP-1 against 26 strains of clinically isolated ESKAPE MDROs (*E. faecium*, *S. aureus*, *K. pneumoniae*, *A. baumannii*, *P. aeruginosa*, and Enterobacter species)^48^ was evaluated and the MICs and minimum bactericidal concentrations (MBCs) were calculated. The MIC experiment showed that QL-AMP-1 exhibited strong growth-inhibitory activity against 20 out of the 26 superbugs (MIC ≤ 64 μg/mL), significantly exceeding LL-37 and offering comparable or superior efficacy to SAAP-148^11^ (an improved AMP derived from human LL-37 through extensive wet lab experiments) against 15 strains of MDROs (Table 1). Typically, an antimicrobial agent is considered bactericidal if its MBC is less than or equal to 4x its MIC^49^. The MBC experiments further confirmed that QL-AMP-1 killed all tested strains except 2 strains at doses not exceeding 4x the MIC, which suggests that QL-AMP-1 is an effective bactericidal agent against MDRO (Table 1).

**Table 1.**
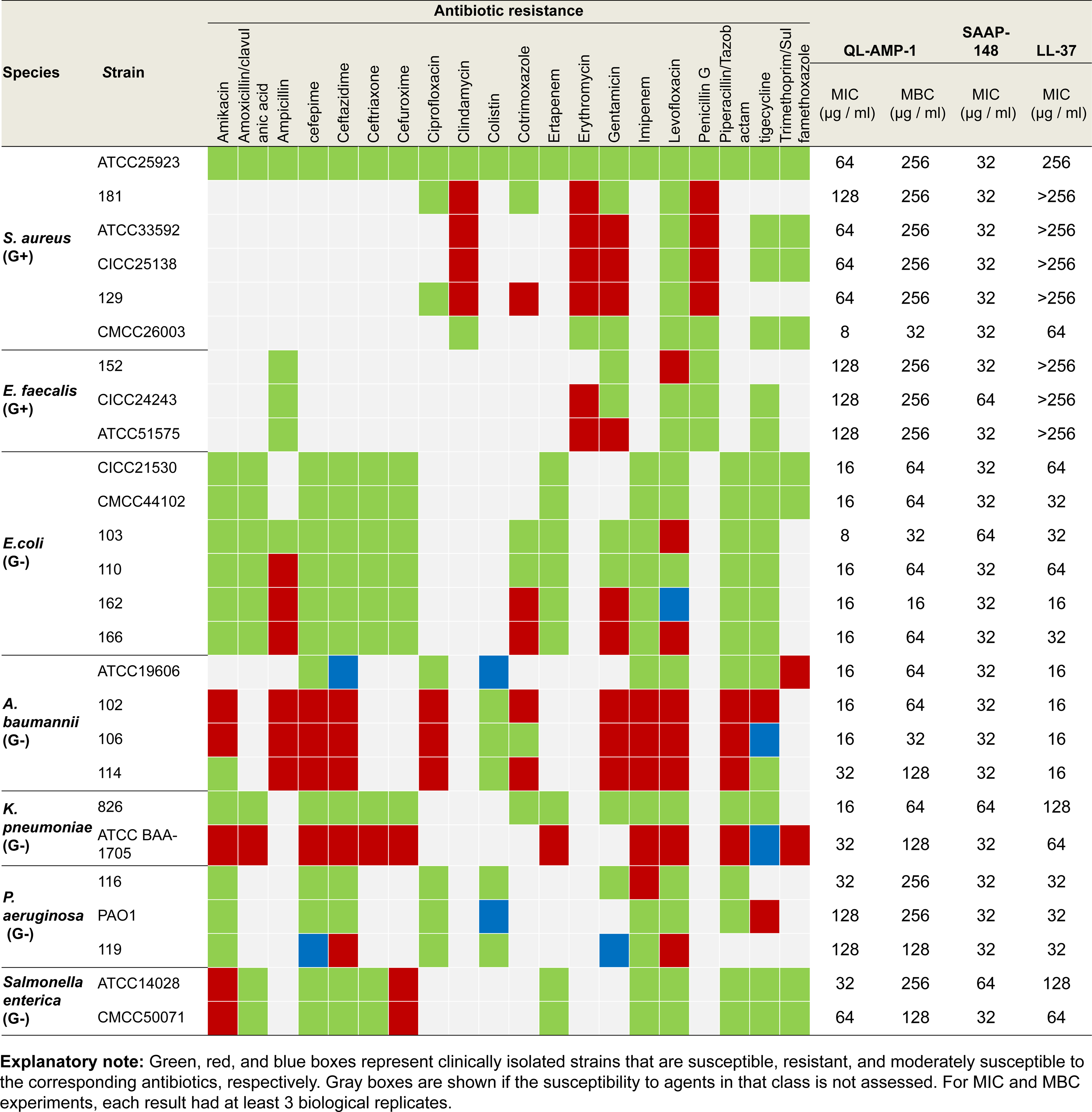
Antimicrobial activities of antimicrobial peptides QL-AMP-1, SAAP-148 and LL-37 against clinically isolated antibiotic-resistant bacteria.

### Off-target toxicity

Off-target toxicity of the AMPs largely determines their potential for clinical application^14^. Cytotoxicity and haemolytic assays to human kidney cell line HEK-293T and human erythrocytes were performed to evaluate the safety of QL-AMP-1 and compared them with those of SAAP-148 (Fig. 4a-d and Supplementary Fig. 3a-d). We calculated the cytotoxic concentrations at which 50% of the HEK-293T cells remained viable (CC50) and the haemolytic concentrations at which 50% of the erythrocytes were lysed (HC50) for QL-AMP-1 and SAAP-148, and computed their therapeutic windows for 26 bacterial strains. The results showed that the HC50 and CC50 of QL-AMP-1 (HC50 = 767.75 μg/mL; CC50 = 234.01 μg/mL) were much larger than that of SAAP-148 (HC50 = 86.08 μg/mL; CC50 = 36.39 μg/mL), and revealed that QL-AMP-1 had a larger safe therapeutic window than SAAP-148 against all 26 strains tested, demonstrating the excellent safety profile of QL-AMP-1 (Fig. 4e and Supplementary Table 9).

**Fig. 4.**
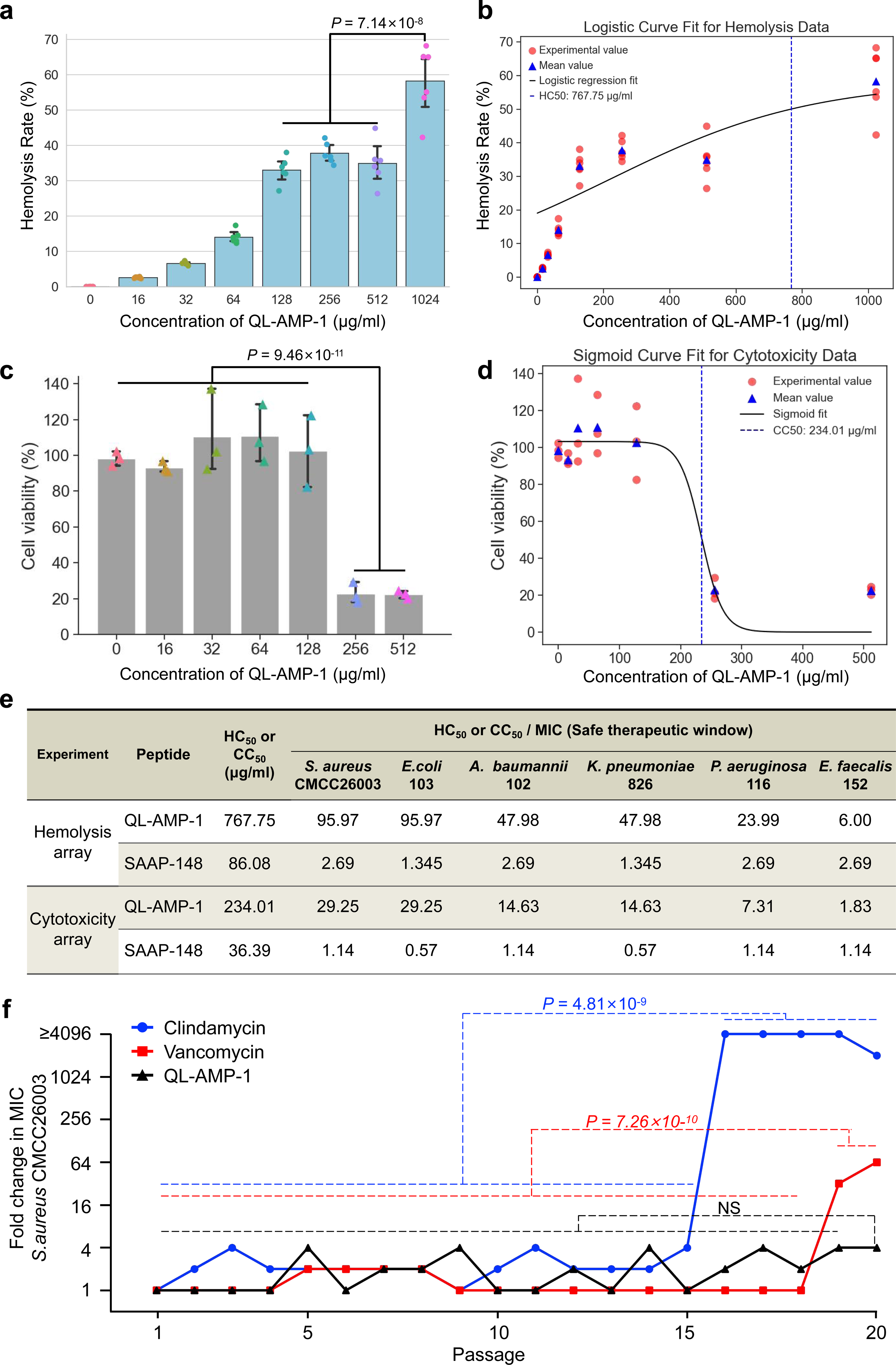
Off-target toxicity and induced drug resistance of QL-AMP-1 *in vitro*. **a-d,** Hemolysis and cytotoxicity of QL-AMP-1 against human red blood cells (a,b) and 293T cells (c,d) at different concentrations. HC50 (b) and CC50 (c) are calculated by logistic regression and sigmoid function respectively (n = 6 biologically independent replicates). **e,** Shown are the antimicrobial activities and safe therapeutic window of AMPs QL-AMP-1 and SAAP-148 against several bacteria. HC50 and CC50 respectively refer to the half hemolytic toxicity concentration of the drug to human red blood cells and the half cytotoxicity concentration to the cell line 293T. **f,** Resistance-acquisition studies of *S. aureus* CMCC26003 when cultured in the presence of sub-MIC (1/2×) levels of clindamycin, vancomycin and QL-AMP-1 (n = 3 biologically independent replicates). All data are mean ± S.D.. NS not significant. All p-values were calculated and reported using two-tailed Student’s t-test.

### Development of microbial resistance

Subsequently, we assessed the ability of *S. aureus* to develop a resistance to QL-AMP-1, using the classical antibiotics clindamycin and vancomycin as references. As shown in Fig. 4f, *S. aureus* cultured at sub-inhibitory concentrations did not exhibit significant resistance to QL-AMP-1 after 20 passages. In contrast, *S. aureus* developed a resistance to clindamycin and vancomycin after 15 and 18 passages, respectively, with their susceptibility decreasing by factors of 8,189 and 64, respectively, by the 20 passages. These results, combined with the data obtained from antibacterial testing against clinically derived MDROs, suggest that QL-AMP-1 holds promise in addressing the rapidly emerging problem of AMR.

### Antimicrobial mechanism

To demonstrate the ability of the M3-CAD pipeline to design novel *de novo* AMPs with multiple bactericidal mechanisms, we examined whether QL-AMP-1 simultaneously possessed four bactericidal mechanisms, namely, bacterial membrane disruption, biofilm formation inhibition, DNA replication suppression, and protein synthesis inhibition.

The mechanism by which most cationic amphipathic AMPs inhibit bacteria is believed to involve electrostatic interactions with bacterial membranes that leads to pore formation and leakage of cell contents^9, 10^. The cationic, amphipathic properties of QL-AMP-1 and the predicted alpha-helical structure suggested bactericidal permeability (Fig. 5a and Supplementary Table 8). Circular dichroism confirmed that QL-AMP-1 exhibited stronger alpha helicity in hydrophobic environments, which simulated the cell membrane (50% TFE), than in aqueous environments (PBS) (Fig. 5b and c, Supplementary Table 10). Confocal fluorescence microscopy of *S. aureus* cells treated with QL-AMP-1 and stained with SYTO 9/PI was used to confirm membrane disruption. SYTO 9 and PI are green and red fluorescent dyes, respectively, which mark cells with intact bacterial membranes and those with membrane pore formation. As shown in Fig. 5d, significant red fluorescence was visible in the group treated with 1×MIC of QL-AMP-1 (64 μg/ml) in contrast to the vehicle, confirming the membrane-disrupting mechanism of QL-AMP-1.

**Fig. 5.**
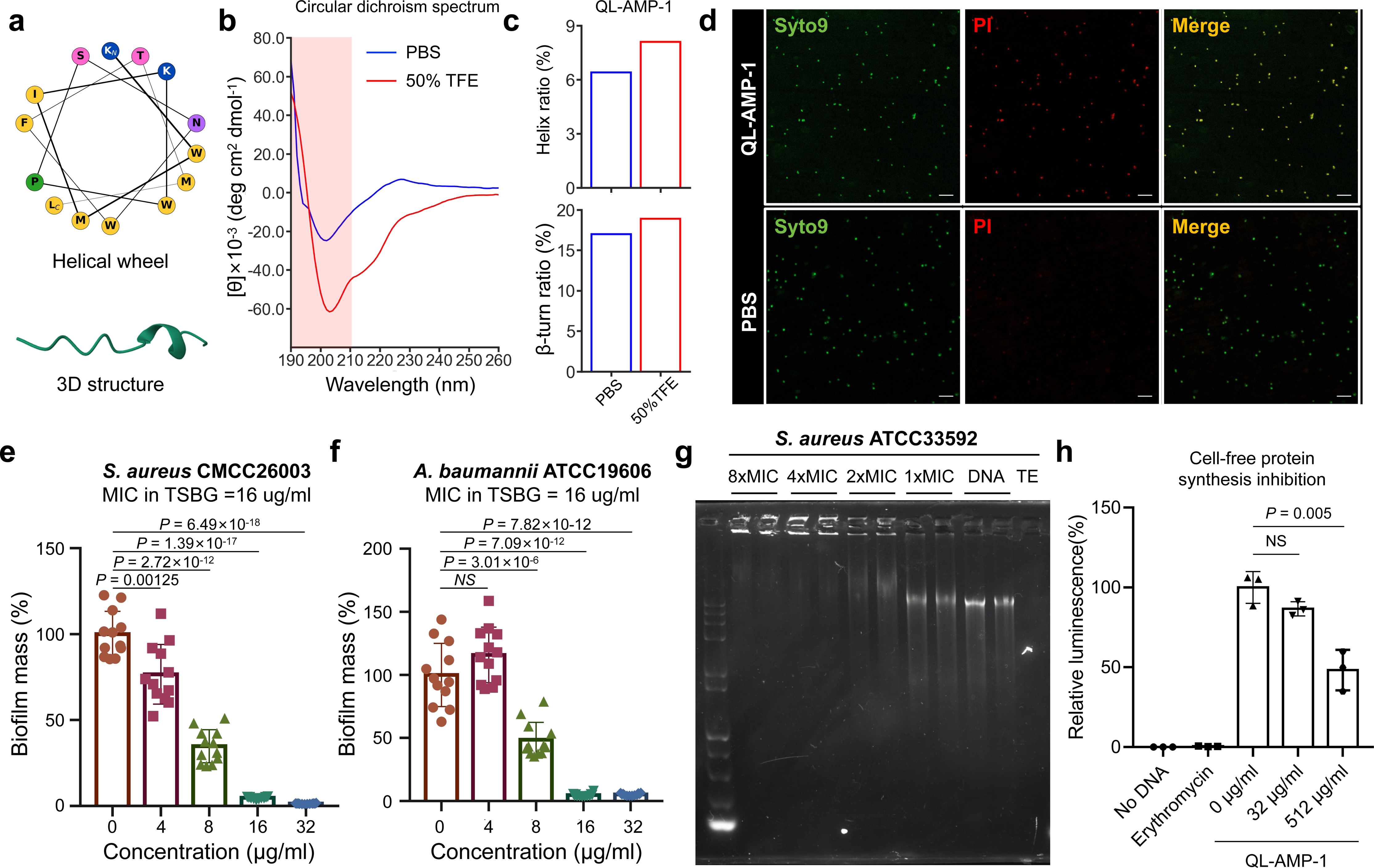
Wet-lab experiments verified the four antimicrobial mechanisms of QL-AMP-1. **a,** Shown are the helical wheel and AlphaFold2 predicted-3D structure diagrams of QL-AMP-1, which illustrates the α-helical structure of the peptide. **b,** CD spectra of QL-AMP-1 in PBS or 50% TFE. The area covered in red indicates the wavelength range of 190-210nm. **c,** Shown are the changes in the helix and β-turn structures of QL-AMP-1 in PBS or 50% TFE solution determined by CD spectra at 190-210nm wavelength. **d,** Confocal fluorescence microscopy of *S. aureus* 33592 cells treated with 64 μg/ml QL-AMP-1 and stained with SYTO 9/PI was used to confirm membrane disruption. SYTO 9 and PI are green and red fluorescent dyes, respectively, which mark cells with intact bacterial membranes and those with membrane poreformation. **e,f,** Shown is biofilm inhibition of *S. aureus* (e) and *A. baumannii* (f) cultured in biofilm-inducing medium (TSBG) with varying concentrations of QL-AMP-1 (n = 12 biologically independent replicates). **g,** DNA binding affinity experiments confirmed that QL-AMP-1 binds to bacterial genomic DNA and blocks DNA migration in agarose gels. **h,** Cell-free protein synthesis inhibition experiments confirmed that the potential of QL-AMP-1 to inhibit bacterial protein synthesis. All data are mean ± S.D.. NS not significant. All p-values were calculated and reported using two-tailed Student’s t-test.

Biofilms enhance bacterial resistance, anti-phagocytic properties, and adherence and are considered a significant factor in the emergence of bacterial AMR^50^. Inhibition of biofilm formation in *S. aureus* and *A. baumannii* were assessed using crystal violet staining. As shown in Fig. 5e and f, QL-AMP-1 inhibited *S. aureus* CMCC26003 and *A. baumannii* ATCC19606 biofilm formation in a dose-dependent manner, and reduced the biofilm mass of both strains by approximately 50% at 0.5×MIC (8 μg/mL). Surprisingly, QL-AMP-1 did not fully inhibit both strains at concentrations of 4 and 8 μg/mL, suggesting that the inhibition of biofilm formation at these concentrations is not due to inhibited bacterial growth.

Next, we examined the potential of QL-AMP-1 to inhibit bacterial DNA replication and transcription through a DNA-binding affinity experiment, in which the binding of AMPs to the bacterial genomic DNA hindered the migration of DNA in agarose gel. As shown in Fig. 5g, QL-AMP-1 started binding to *S. aureus* ATCC33592 genomic DNA at 2×MIC, fully binding at greater than or equal to 4×MIC, suggesting DNA replication and transcription inhibition.

Finally, cell-free protein synthesis inhibition assays demonstrated the impact of QL-AMP-1 on Renilla luciferase protein expression in *E. coli* at 512 μg/mL (Fig. 5h). These results suggested that QL-AMP-1 exhibited multiple bactericidal mechanisms.

### *In vivo* therapeutic efficacy of QL-AMP-1

Potential adverse reactions to local application of QL-AMP-1 in mice were evaluated (Supplementary Fig. 4a). Treatment of both shaved intact and abraded skin with 5× the therapeutic dose (1.5 mg) of QL-AMP-1 showed no primary irritation at 72 h post-treatment. No significant behavioural or body-weight changes were observed, and no signs of systemic toxicity were detected. Histological examination revealed histopathological findings comparable to those of the untreated, control-treated, and QL-AMP-1 solution-treated skin (Fig. 6a-c and Supplementary Fig. 4b-d).

**Fig. 6.**
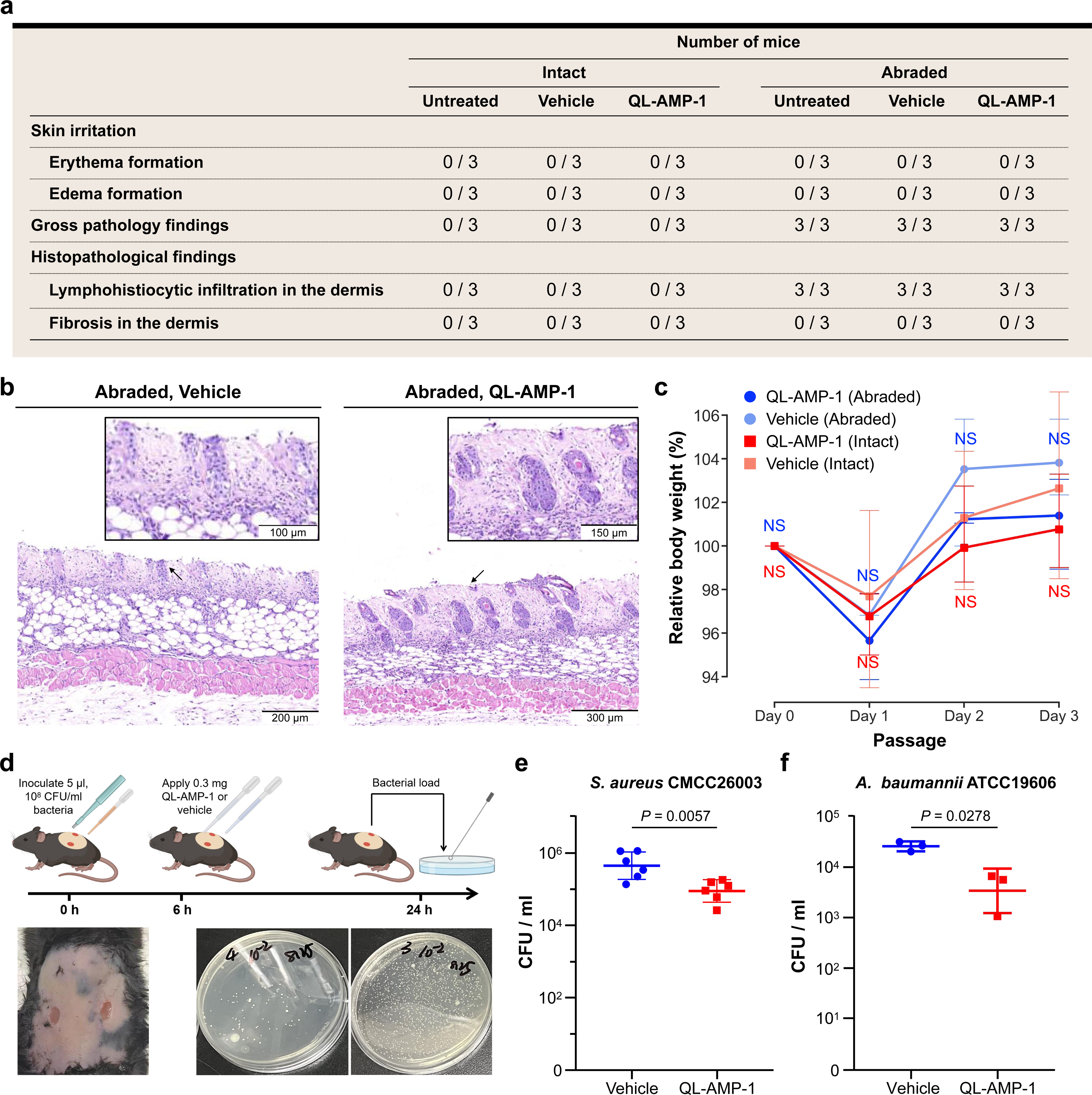
Safety and therapeutic efficacy of QL-AMP-1 in treating skin infections. **a,** Mouse skin tolerance test. Mice with intact or abraded skin were treated with 5× the therapeutic dose (1.5 mg) of QL-AMP-1 or vehicle. As controls, untreated mice were included. Results are expressed as the number of animals out of the total number of animals within the group (n = 3) that showed signs of skin irritation or pathology within 72 hours after treatment. **b,** Representative microscopic imaging of tissue biopsies of abraded skin treated with vehicle or QL-AMP-1 (n = 3 biologically independent replicates). **c,** Body weight changes in mice treated with QL-AMP-1 or vehicle with intact or abraded skin (n = 3 biologically independent replicates). **d,** Schematic representation of bacterial infection and QL-AMP-1 treatment experiments in a full-thickness skin wound model. Created with BioRender.com. **e,f,** Shown is the effect of therapeutic doses of QL-AMP-1 (0.3mg) or vehicle on tissue bacterial burden in a full-thickness skin wound model of *S. aureus* (e, n = 6) or *A. baumannii* (f, n = 3) infection. All data are mean ± S.D.. NS not significant. All p-values were calculated and reported using two-tailed Student’s t-test.

Next, we evaluated the *in vivo* therapeutic effects of QL-AMP-1 in a model of full-thickness skin wounds in mice which were locally infected with *S. aureus* or *A. baumannii*. As shown in Fig. 6d, after removing the hair from the back of the mouse, two full-thickness skin pieces with a diameter of 5 mm were removed from the centre of the back using a tissue biopsy device to establish a full-thickness skin wound model. Each of the two wounds on each mouse was injected with 5 ul of 108 CFU/mL bacterial solution. After 6 hours, the treatment group was given 0.3 mg QL-AMP-1. 24 hours post-modelling, the skin from the infected area of the animal was aseptically collected, and live bacteria were detected and counted. The results revealed that, compared to the vehicle, QL-AMP-1 significantly reduced the bacterial load of *S. aureus* and *A. baumannii* in the full-thickness skin wound model by approximately 79.96% and 86.77%, respectively (Fig. 6e and f).

## DISCUSSION

Remarkable progress has been made in the discovery of AI-driven AMPs in recent years^16–28^. Regardless of the task (that is, identification, optimisation, or generation) and the underlying network used, existing AI models inherently link the primary sequence of peptides and/or their sequence-derived physicochemical properties to their bactericidal activity. Consequently, the information scope provided by the primary sequence inherently caps the potential of all methods that utilise these features for modelling. In this study, we sought to address a critical scientific query. Could there be undiscovered features and functional properties that could bolster the AI-driven discovery of novel AMPs beyond the conventional sequence features and antimicrobial activity properties?

By leveraging computational methods such as AlphaFold2^30^ and RoseTTAFold^31^, we are now capable of predicting 3D protein structures with near-experimental accuracy and atomic-level precision. We postulated that integrating the 3D structural features of peptides could enhance the performance of the AI in AMP design. Ablation studies supported our multimodal model, which combines peptide primary sequence features with AlphaFold2-predicted 3D structural features, and outperformed models relying solely on primary sequences or 3D structures in multilabel classification tasks concerning AMPs’ antimicrobial activity, inhibition mechanisms, and host toxicity. As for the Functional properties of AMPs, we hypothesised that AMPs featuring multiple antimicrobial mechanisms may exhibit enhanced bacterial inhibition efficacy, particularly against resistant superbugs. To investigate this, we reviewed the literature and manually annotated four known inhibitory mechanisms across 12,914 AMPs that served as training data for the M3-CAD pipeline, and enabled us to design and screen AMPs that embody all four antimicrobial mechanisms. Moreover, we integrated variations in the inhibitory activities of the AMPs against different bacterial types into the model, hypothesising that this could facilitate broad-spectrum AMP design. Wet lab evidence on QL-AMP-1 confirmed that the M3-CAD pipeline, trained with AMPs’ antimicrobial mechanism and inhibitory activity data against six types of resistant bacteria, was able to generate multi-mechanism AMPs that inhibited both gram-positive and gram-negative MDROs.

During the construction and training of the M3-CAD pipeline, two critical technical queries were confronted: the representation of polypeptide 3D structures with a long-tail length distribution and the imbalance of functional labels of AMPs in multilabel classification tasks. For the first query, GNNs were proposed for representing the 3D structures of proteins predicted by AlphaFold2. The dimensions of the GNN’s adjacency matrices equalled the number of amino acids in the longest protein in the training set^51^. However, GNNs seem unsuitable for representing polypeptide 3D structures in AMP discovery because the lengths of the available AMPs for training show a long-tail distribution. With an increase in sequence length, the number of polypeptides continued to decrease, which led to an overly sparse GNN adjacency matrix. Moreover, because of the sequential connection of amino acids within a polypeptide, relying on a single path for feature transformation may hinder feature transmission between distant amino acids. Therefore, we proposed a 3D voxel-coloring method that transforms the problem into a 3D visual feature extraction task, addressed using a 3D-CNN model that outperformed traditional GNNs for AMP discovery tasks. Regarding the second query, we introduced a multilabel rebalancing loss function to improve the performance in all three multilabel tasks.

The speed and accuracy of M3-CAD in AMP discovery are noteworthy. In a single run, it can generate and rank 200,000 peptides in approximately two hours. Wet lab validation confirmed *in vitro* antimicrobial capabilities of all ten high-priority peptides. QL-AMP-1, the most promising candidate, exhibited multiple antimicrobial mechanisms, broad-spectrum activity against MDROs, low host toxicity, and reduced resistance-inducing propensity. With further modifications, such as N-terminal acetylation or D-amino acid substitution^14, 52^ it holds promise for advancing clinical trials, thereby highlighting the potential of M3-CAD to expedite future AMP research and translational efforts.

To summarize, from the perspective of AI-driven AMP design problems, we have demonstrated that the introduction of polypeptide 3D structures led to a significant performance boost in multimodal models beyond models that solely utilise primary sequences or 3D structures. The proposed 3D voxel-coloring method aids in enhancing the representation of polypeptide 3D structural features, which further improved the downstream task performance. Including antimicrobial mechanism information and differential inhibitory activity data of AMPs against various bacteria as training data enabled the AI to discover AMPs with multiple antimicrobial mechanisms and broad-spectrum antimicrobial activity. These discoveries will foster future automated AMP design research with potential generalisable methodological implications for AI-driven therapeutic peptide discovery.

## METHODS

### Collection and annotation of training data

We compiled an AMP database, named QLAPD, which encompasses 12,914 AMP and 22,411 non-AMP entries. Each AMP recording its amino acid sequence, 3D structure, and six functional attributes, including: (1) Inhibitory activity against six drug resistant bacteria, including *E. faecium*, *S. aureus*, *K. pneumoniae*, *A. baumannii*, *P. aeruginosa*, and Enterobacter; (2) Number of types of non-resistant bacteria it can inhibit (ranging from 0 to 6); (3) Number of types of drug resistant bacteria it can inhibit (ranging from 0 to 6); (4) Four antimicrobial mechanisms: lipid bilayer disruption, cytoplasmic protein inhibition, DNA/RNA binding, and biofilm inhibition; (5) Toxicity; (6) Six kinds of organ toxicity: cardiotoxin, enterotoxin, neurotoxin, hemostasis impairing toxin, myotoxin, and others. Below we provide details of the functional attributes and 3D structure feature:

Details of AMP/non-AMP annotation: The sequences of AMPs were primarily obtained from DBAASP^32^, dbAMP^33^, and DRAMP^34^ repositories, with a few sourced from literature and patent databases such as PubMed and Google Scholar. We manually screened and binarized the inhibitory activity data of each AMP against six types of bacteria: *E. faecium*, *S. aureus*, *K. pneumoniae*, *A. baumannii*, *P. aeruginosa*, and Enterobacter. Specifically, if an AMP has a MIC of ≤ 256 μg/ml against any resistant strain of a certain type of bacteria, the AMP is considered active against that resistant bacteria type, and vice versa. The same principle applies to non-resistant strains. The non-AMP instances with lengths of 10-50 amino acid residues were collected from the UniPort database^44^, after discarding sequences that were annotated as AMP, membrane, toxic, secretory, defensive, antibiotic, anticancer, antiviral, and antifungal and were used in this study as well.

Details of toxic/non-toxic annotation: Sequences with toxicity labels were curated from DBAASP^32^ database. Sequences with IC50/HC50 ≤ 256 µg/ml in DBAASP were considered as Toxic instances. Sequences of 10-50 amino acid residues in the Uniport database were defined as nontoxic after excluding sequences that have been defined as toxic in the DBAASP and UniPort databases. For other mechanism-based toxic, we use the label from the uniport database. For the organ-based toxic, if an AMP has the toxin to one type of the organ, then we regard the AMP has the organ toxicity.

Details of antimicrobial mechanism annotation: We annotated the lipid bilayer disruption, cytoplasmic protein inhibition, DNA/RNA binding, and biofilm inhibition mechanisms of the AMP by directly using the properties from the DBAASP database^32^.

Details of peptide’s 3D structure: The 3D structure of the peptide was predicted by the AlphaFold2 model^30^, and stored in the format of the PDB^36^, which records the type of each atom in the peptide, the coordinates of the atom in 3D space, and the type of amino acid residue to which the atom belongs.

### The 3D voxel coloring method

Based on the three-dimensional coordinates, van der Waals radii of each atom, atomic mass, amino acid solubility, amino acid equipotential, and other known information, we constructed a multi-channel polypeptide structure coloring method. Specifically, the method uses the spatial structure of the atoms in the polypeptide sequence to determine the space occupied by the polypeptide. Taking the center of gravity of the polypeptide as the reference point, the x-axis, y-axis, and z-axis are constructed, with a unit distance of 1 angstrom on each axis. The method treats this unit distance as a voxel in a 3D grid. It is assumed that only atoms within a range of L angstroms in the positive and negative directions of each coordinate axis are considered. The entire 3D voxel is represented as a cube with dimensions L*2 in length, width, and height, where L is any positive integer and is typically set as a multiple of 8 to facilitate subsequent feature extraction. The van der Waals radius of each atom, rounded to angstroms (with the rounding for each atom as follows: ’H’: 1, ’C’: 1.5, ’N’: 1.5, ’O’: 1.5, ’S’: 2)^37^, determines the space it occupies in the cube.

To leverage the inherent properties of atoms and various amino acids, the method innovatively presents a multi-channel polypeptide coloring approach inspired by the RGB multi-channel visual imaging concept. Specifically, the properties of the atom (atomic mass), the solubility of the amino acid it constitutes, and the amino acid’s acidity or basicity are treated as three distinct channels for coloring the peptides. In the first channel, the atomic mass is rounded and used directly as the voxel value. In the second and third channels, related to amino acid properties, the spatial positions of multiple atoms can represent the position of a single amino acid. Therefore, for the second solubility channel, the solubility of different amino acids is categorized into hydrophobic and hydrophilic amino acids. To differentiate these two types, and with a background voxel value of 0, the method assigns a value of 128 to hydrophobic amino acids and a value of 255 to each atom of hydrophilic amino acids. Similarly, for the third pH channel, the voxel values for acidic, neutral, and basic amino acids are assigned as 86, 168, and 255, respectively.

### Multi-modal conditional varitional auto-encoder for peptide generation

The variational auto-encoder has been a powerful tool to generate the anitbiotic peptide recently. It is mainly based on learning the meaningful lantent data embedding from the training data with a reconstruction loss and a regulation loss. The training process on sampling the data could be formulated as:

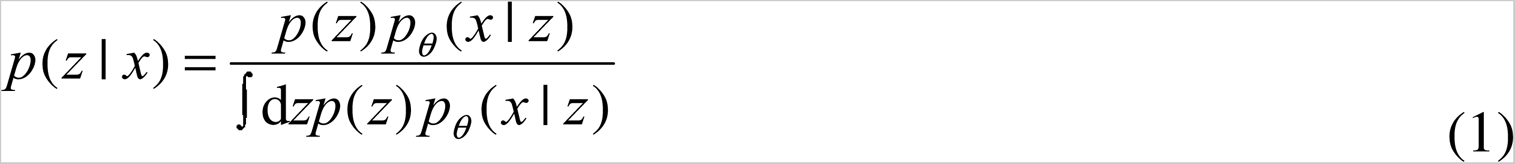

To enable the conditional generation of the target data, conditional VAE was proposed with an additional category supervision loss term, enables the feature embedding could be directly applied to other situations. This process could be formulated as follows:

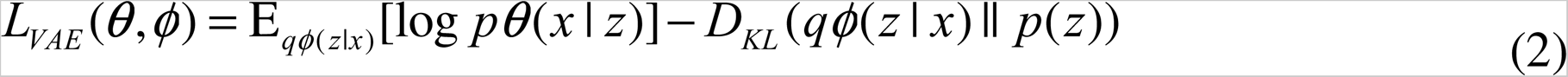

However, all the above mentioned autoencodes only task the toy cases into account. The ignorance of the structure could do harm to the learning process of the peptide generation. Moreover, there is another question on how to embed the condition information into the training process of the multi-modal cVAE. Thus, in this work we propose use a novel Unlike the previous VAE that only takes the sequence as the input, our multi-modal cVAE has two encoders and two decoders, which are used to encode/decode structure input and sequence input, respectively. In order to generate the AMPs with the excepted attributes, we need to inject the conditional information (e.g., broad-spectrum drug-resistant, non-toxin) into the autoencoder.

### Multi-label multi-modal rebalancing learning for attribute identification

To further identify the MDR AMPs from the generated sequences, we proposed a efficient multi-modal multi-label model to take advantage of both sequence information and structure information. Specifically, we use a 3D-ResNet *f_struct_* to capture the structure information *x_struct_* based on the abovementioned structure coloring features. To extract the sequence feature *x_seq_* of AMPs, we use a multi-layer neural network *f_seq_* to capture its feature. After that, the extracted high-level features are concatenated and sent to a classification head for multi-label classification. The classification head *f_head_* is a multi-layer neural network that takes the fused feature as the input and the label number of the target task as the output. Let *y* be the prediction result in the form of logiest function, this process is formulated as:

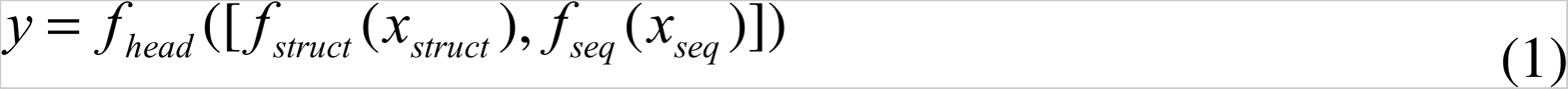

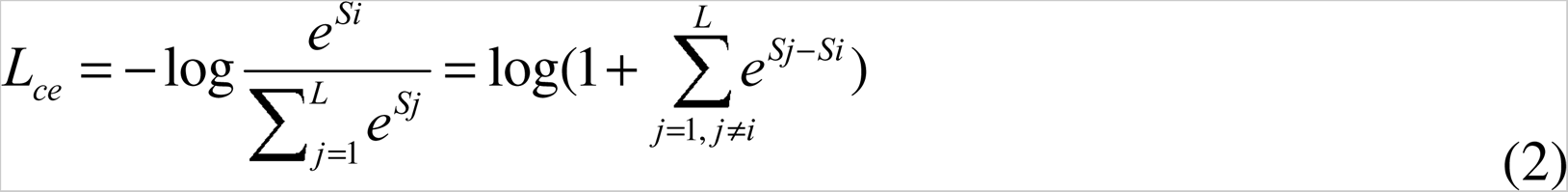

Considering that the imbalance data distribution will hard the learning process of the deep neural network model, we adopt a re-balancing loss to handle the imbalanced multi-label classification task. Unlike unbalanced multi-class classification that can simply increase the number of the samples on the minority categories, in multi-label classification task, increase the number of the minority category may lead to worse imbalance as the samples within minor category may also have the majority category. The above mentioned phenomenon motivates us to propose a re-balancing loss for the unbalanced multi-label classification task. Specifically, we extend the “softmax + cross entropy” scheme to a multi-label classification scene with multiple target classes. The method is based on the consideration that “the score of each target class is not less than that of each non-target class”. The balance loss function is obtained to handle the imbalance data distribution.

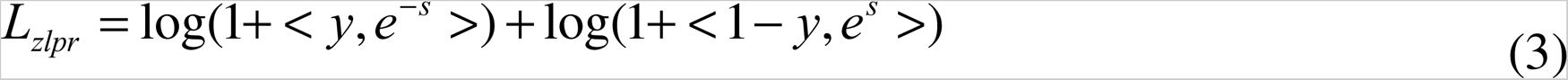

### Method for calculating the physicochemical properties and sequence similarity of the peptides

Physicochemical properties such as hydrophobicity, hydrophobic moment, aliphatic index, aromaticity, instability index, alpha-helix, beta-sheet, turns, charge, isoelectric point, charge density, and flexibility were estimated using the ProteinAnalysis function of the BioPython library^53^.

The similarity between sequences was calculated using the Needleman-Wunsch algorithm, a fundamental method for sequence alignment. This algorithm is implemented in the BioPython library^53^, specifically through the pairwise2 module. A function named calculate_similarity was created to facilitate this task. It accepts two sequences in string format, seq_gen and seq_tem, and converts them into Seq objects, as defined in BioPython. The sequences are then globally aligned using the pairwise2.align.globalms method, which stands for "global alignment using the Needleman-Wunsch algorithm with custom scoring." The scoring system includes a match score of 1, a mismatch score of -1, a gap opening penalty of -0.5, and a gap extension penalty of -0.1. The function returns the score of the optimal alignment divided by the length of the first sequence (seq_gen). This normalized score provides a measure of similarity between the two sequences, taking into account differences in their lengths.

### Strains and Animals

Standard strains included *E. faecalis* CICC24243, *E. faecalis* ATCC51575, *S. aureus* ATCC25923, *S. aureus* ATCC33592, *A. baumannii* ATCC19606, *Salmonella enterica* ATCC14028, *E.coli* CICC21530, *S. aureus* CICC25138, *K. pneumoniae* ATCC BAA-1705 were purchased from China Center of Industrial Culture Collection (CICC). Standard strains included *E.coli* CMCC44102, *Salmonella enterica* CMCC50071, *S. aureus* CMCC26003, *P. aeruginosa* PAO1 were purchased from National Center for Medical Culture Collections (CMCC). Standard strains included *S. aureus* ATCC25923 were purchased from Ningbo Mingzhou Biotechnology Co., LTD. The clinical multidrug-resistant strains including *K. pneumoniae* 826, *E.coli* 103, *E.coli* 110, *P. aeruginosa* 116, *E. faecalis* 152, *S. aureus* 181, *S. aureus* 129, *A. baumannii* 102, *A. baumannii* 106, *A. baumannii* 114, *E.coli* 162, *E.coli* 166, *P. aeruginosa* 119 were obtained from the clinical Laboratory of Qilu Hospital. The detection of bacterial resistance to various antibiotics was completed by the clinical Laboratory of Qilu Hospital. All strains were streaked on Nutrient Broth (NB) agar medium and incubated at 37 °C overnight.

Female C57BL/6J mice were purchased from Beijing Vital River Laboratory Animal Technology Co., Ltd. They were cultured within sterile isolators at the Laboratory Animal Centre of Shandong University on a 12 h - 12 h light-dark cycle with ambient temperature of 20-26°C, humidity of 40 - 70%, and ad libitum access to food and water. All the animal experiments involved were approved by the Animal Care and Animal Experiments Committee of Qilu Hospital of Shandong University (DWLL-2023-104).

### Antimicrobial activity assays

Antimicrobial activity of antimicrobial peptides was examined in sterile 96-well plates with the broth microdilution according to CLSI standard. Suspended the colonies in saline solution, adjust the turbidity to McFarland 0.5 to reach the bacteria concentration of 10^8 CFU/ml and then again diluted 100 times for the inhibition test. 50 μL of bacterial suspension at 1×10^6^ CFU/mL were incubated with the same volume of different concentrations of peptide solution [serial 2-fold dilutions in Mueller-Hinton Broth (Hopebio)]. After incubation for 18–20 h at 37 °C, the MIC that no obvious bacteria could be observed was recorded. 50 μL of the mixture without observed bacteria was added to MH-agar plates. After incubation for 18 h at 37 °C, the MBC without bacteria growth was recorded. All experiments were performed with three independent replicates.

### Analysis of cytotoxicity

Human embryonic kidney (HEK-293T) cells were obtained from the American Type Culture Collection (ATCC) and grown 37 °C in a humidified atmosphere containing 5% CO2 in Dulbecco’s modified Eagle’s medium (DMEM) supplemented with 1% antibiotics (penicillin and streptomycin) and 10% fetal bovine serum (FBS). Cytotoxicity of peptides against 293T cells was determined using the CCK8 assay. HEK-293T cells were seeded on 96-well microplates at a density of 8×10^3^ cells/well in 100 ul above culture medium and cultured at 37L for 24h. Then the culture medium was removed and 100 ul culture medium with peptide at the serial 2-fold dilutions concentrations were added to each well. Wells containing cells without peptides served as controls. The cells were cultured for another 24 h, followed by replacement of the media with 10% CCK-8 containing fresh DMEM medium and another 1 h incubation at 37 °C. The absorbance of the solution was measured at 450 nm by a microplate reader. The cell viability was calculated according to the equation:

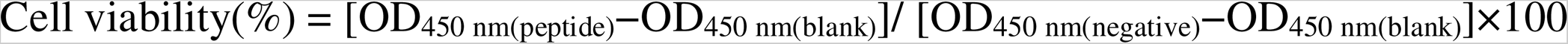

For haemolysis test, fresh human red blood cells (hRBCs) were washed 3 times with PBS (35 mM phosphate buffer, 0.15M NaCl, pH 7.4) by centrifu-gation for 5 min at 1000 × g, and resuspended in PBS. 50 ul peptide solutions (serial 2-fold dilutions in PBS) were added to 100 ul of hRBC suspension [8%(v/v)in final] in PBS and incubated for 1 h at 37 L. Samples were centrifuged at 1000 × g for 5 min, and hemoglobin release was monitored by measuring the supernatant absorbance at 570 nm with a flat bottom 96 wells plate (servicebio). hRBCs in PBS or 0.1% Triton X-100 were used as the negative and positive controls, respectively. The hemolysis percentage was calculated according to the equation:

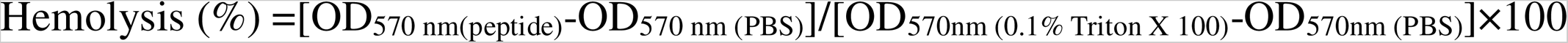

We estimated by non-linear regression the peptide dose that led to 50% hemolysis (HC50) and 50% cytotoxicity (CC50)

### CD Spectroscopy

CD spectroscopy was used to study the secondary structure of peptides. The peptide was dissolved at 50uM in either 10 mM PBS (pH 7.4) or 50% trifluoroethanol in PBS. TFE is cosolvent known to stabilize secondary structures. Added peptide solutions to the cuvette. The optical path of the CD spectra was 1 mm, the scan wavelength range was 190 to 260 nm, and the scan rate was 1nm/s. The mean residue ellipticity was calculated using the following equation:

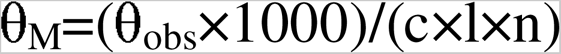

where θ_M_ is the residue ellipticity (deg cm^2^ dmol^−1^), θ_obs_ is the measured ellipticity (mdeg), c is the peptide concentration (mM), l is the path length (mm), and n is the number of amino acids.

### Drug resistance assay

For comparison, development of resistance to the clinically relevant antibiotics clindamycin (Macklin) and vancomycin (Macklin) was determined. The evolution of resistance by *S. aureus* CMCC26003 to either antibiotics or peptide was monitored for 20 days of serial passaging in liquid NB. Cells were passaged every 24 h. Briefly, 1ml of bacterial suspension (1×10^6^ CFU/mL) were incubated with antibiotics or peptide solution at 1/2 minimal inhibitory concentration at 37L, 200 rpm. After each incubation period, a new inoculum of 10^6^ cells was prepared for inoculation of the following passage containing fresh medium and increased doses of the antimicrobial agent. During the 20 days of serial passaging, the minimum inhibitory concentration of antibiotics or peptide to each generation of cells was determined. MIC determination method was as previously described. The development of bacterial resistance was defined as a more than fourfold increase in MICs compared to the initial MICs.

### The mechanism experiments of AMPs

Membrane permeabilization assay. The membrane permeability of the peptide was determined by using the LIVE/DEAD BacLight Bacterial Viability Kit (Invitrogen). *S. aureus* ATCC33592 were grown to mid-log-phase, centrifuged (5000 × g for 10 min), washed and re-suspended in PBS to 1×10^6^ CFU/ml. The peptide solution was added to 100ml of this bacterial suspension so that the final concentration of peptide was equal to the minimum inhibitory concentration. bacterial cells were incubated with peptides at room temperature for 120 min. Bacterial cells in PBS were used as the negative controls. Then, they were centrifuged at 5000 × g for 5 min and resuspend in 50 μl staining solution. Mix thoroughly and incubate at room temperature in the dark for 15 minutes. Trap 5 μL of the stained bacterial suspension between a slide and an 18 mm square coverslip. Observe in confocal microscopy (LSM980) equipped with specified filter sets.

Anti-biofilm formation. The microtiter dish biofilm formation assay was used to detect the preventive effect of peptides on biofilm formation. TSBG [tryptic soy broth supplemented with 1% (wt/vol) glucose] wass used for the growth of bacterial biofilms. In order to prevent the MIC values measured in different media are inconsistent, the MIC of *S. aureus* CMCC26003 and *A. baumannii* ATCC19606 in TSBG was retested and the result was both 16 μg/mL. The method was improved on the basis of MIC detection. 50 μL of bacterial suspension at 1×10^6^ CFU/mL were incubated with the same volume of peptide solution ranging from 4 to 32 μg/mL in 96-well microplates. As an untreated control, bacteria were exposed to TSBG medium without peptide. After 24 hours incubation at 37L, planktonic bacteria were removed. Gently wash the wells 3 times with PBS, then dry at 37 L. Biofilms were stained with 0.3% crystal violet (Sigma-Aldrich) for 15 min, washed, and solubilized with 30% acetic acid in water. Quantify absorbance in a plate reader at 550 nm using 30% acetic acid in water as the blank. Biofilm mass was calculated according to the equation:

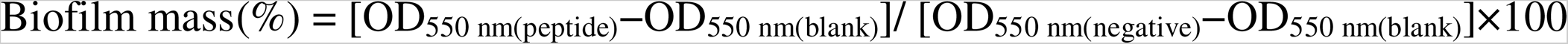

DNA binding assay. The DNA binding ability of peptides was investigated by gel retardation assay. The genomic DNA of *S.aureus* ATCC33592 was extracted by TIANamp Bacteria DNA Kit. 10 ul DNA (30ng/ul) was incubated with the same volume of different concentrations of peptide solution (1× to 8× MIC in final) dissolve in TE buffer for 30 min. As an untreated control, DNA was exposed to TE buffer without peptide. Subsequently, 4 ul of loading buffer was added, and a 20 ul aliquot subjected to 1% agarose gel. Finally, the migration of DNA bands was observed by ultraviolet (UV) illumination with the ImageQuant 300 gel documentation system.

Cell-free protein synthesis inhibition assay. The reactions were performed using the S30 T7 High-Yield Protein Expression System (Promega). 3 μl of peptide solution were added to reaction mixtures to get a final concentration respectively of 32 μg/ml or 512 μg/ml in a final volume of 15 μL in nuclease-free PCR tubes. Nuclease-free water was added to the reaction instead of the peptides as untreated control. Nuclease-free water was added to the reaction instead of peptides and DNA template as blank control. Erythromycin was used as positive control. The samples were incubated for 1 h at 37L with vigorous mixing (300 rpm), and the reaction was stopped transferring the samples on ice. The activity of the Renilla reniformis luciferase, used as a reporter, was quantified using the commercial kit Renilla-Glo Luciferase Assay System (Promega). Mixing 2.5 μl of reaction mixtures and 97.5 μl of the provided buffer, and waiting 10 min before the measurement. Black flat-bottom 96-well plates were used in a luminometer Plate Camaleon (Bioteck). In all the luminescence measurements, the relative values were calculated as a percentage of the untreated control.

### *In vivo* experiments

The therapeutic potential of QL-AMP-1 *in vivo* was tested in a local *S. aureus* CMCC26003 or *A. baumannii* ATCC19606-infected full-thickness skin wound model. Ten-week-old female C57BL/6J mice were anesthetized using isoflurane and administered buprenorphine as an analgesic at 0.1 mg kg−1 intraperitoneally. Then, we removed the back hair of mice, two pieces of full-layer skin with a diameter of 5 mm were removed from the center of the back of mice with a tissue biopsy device. The model was successfully established if the wound area of mice in each group was uniform. Mice were immediately infected with 5 uL of 10^8^ CFU/mL bacterial solution directly pipetted onto the wound bed. The wound is covered with sterile dressing. After 6 h, the treatment and control group were given 10 uL of 30 mg/ml antimicrobial peptide QL-AMP-1 or 10 uL H_2_O, respectively. Mice were killed at experimental endpoint (24 h postinfection) and wound tissue was collected, homogenized in phosphate-buffered saline and plated on solid LB medium. Count the number of colonies on LB medium the next day.

### Safety

For the safety study, the intact or abraded skin of the mice was treated once with antimicrobial peptides QL-AMP-1 or water. In brief, ten-week-old female C57BL/6J mice were anesthetized using isoflurane and administered buprenorphine as an analgesic at 0.1 mg kg−1 intraperitoneally. After removing the back hair of the mice, the skin on the back of the mice in the scratch group was snded with 60-grit sandpaper until the skin oozed blood. The intact skin group received no additional treatment except shaving. 50 ul QL-AMP-1 (30 mg/ml) or water was applied uniformly over the intact or abraded skin to assess the acute dermal toxicity. Groups without any addition served as untreated controls. Histopathological characteristics of treated skin detected by H&E staining on the third days.

## ACKNOWLEDGMENTS

We thank the Translational Medicine Core Facility of Shandong University for consulting and lending instrument that supported this work. The scientific calculations in this study were performed on the HPC Cloud Platform of Shandong University. This work was financially supported by the National Natural Science Foundation of China (Grant No. 82070552 and 82270580), National Clinical Research Center for Digestive Diseases supporting technology project (Grant No. 2015BAI13B07), and the Taishan Scholars Program of Shandong Province.

## AUTHOR CONTRIBUTIONS

Y.L. initiated, designed, and supervised the study. Y.W., H.G., B.W. and Y.Z. designed and implemented the algorithm. X.L., L.L., and Y.Z. designed and performed the wet laboratory experiments. Y.W., H.G., X.L., L.L., Y.Z., P.B., Q.K., J.N., Z.H., X.N., K.J. and L.S. collected and annotated the data. Y.W., H.G., and X.L. contributed to data visualisation. All authors contributed to the interpretation of the results. Y.W. wrote the original manuscript. Y.W., H.G., X.L., and Y.L. reviewed and edited the manuscript.

## COMPETING INTERESTS

The authors have filed the following patent applications for this work: (1) Application No. CN 116153435A for the method for structural characterisation of peptides based on 3D voxel coloring (inventors: Y.W., Y.L., H.G., L.L., X.L., and X.Z.); (2) Application No. CN 116206690A for the method and system for generating and identifying antimicrobial peptides (inventors: Y.L., Y.W., H.G., X.L., L.L., and X.Z.).

