## Supplementary Figures for "De novo multi-mechanism antimicrobial peptide design via multimodal deep learning"

### Supplementary Information

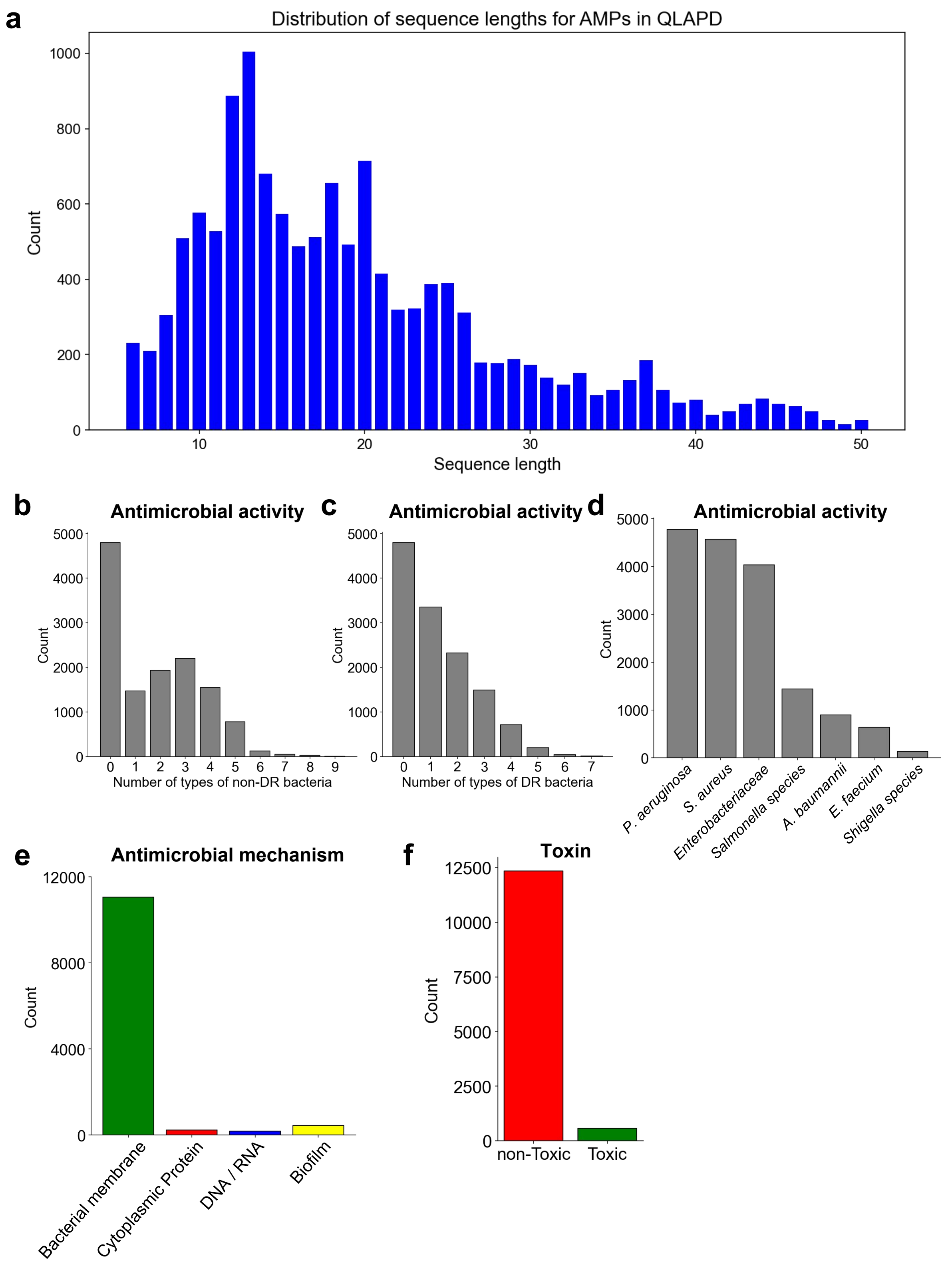

**Supplementary Fig. 1 | Distribution of characteristics and functional attributes of AMPs in the QLAPD database. a,** Distribution of sequence lengths for AMPs (n = 12,914). **b,c,** The distribution of the number of types of non-DR (b) and DR (c) bacteria that AMPs can inhibit. **d,** Distribution of bacterial species that AMPs can inhibit. **e,** The distribution of types of antimicrobial mechanisms of AMPs. **f,** The distribution of toxic and non-toxic AMPs.

**Supplementary Table 1**. Normalized values of atomic features and amino acid features for the 3D voxel coloring method.

| | Atomic name | Abbreviation | Van der Waals radii | Relative atomic mass |
|---|---|---|---|---|
| Atomic features | Hydrogen | H | 1 | 1 |
| | Carbon | C | 1.5 | 12 |
| | Nitrogen | N | 1.5 | 14 |
| | Oxygen | O | 1.5 | 16 |
| | Sulfur | S | 2 | 30 |
| | **Amino acid name** | **Abbreviation** | **Solubility** | **Charge** |
| Amino acid features | Alanine | A | 255 | 155 |
| | Valine | V | 255 | 155 |
| | Proline | P | 255 | 155 |
| | Phenylalanine | F | 255 | 155 |
| | Tryptophan | W | 255 | 155 |
| | Isoleucine | I | 255 | 155 |
| | Leucine | L | 255 | 155 |
| | Glycine | G | 155 | 155 |
| | Methionine | M | 155 | 155 |
| | Tyrosine | Y | 55 | 155 |
| | Serine | S | 55 | 155 |
| | Threonine | T | 55 | 155 |
| | Cysteine | C | 55 | 155 |
| | Asparagine | N | 55 | 155 |
| | Glutamine | Q | 55 | 155 |
| | Aspartic acid | D | 55 | 55 |
| | Glutamate | E | 55 | 55 |
| | Lysine | K | 55 | 255 |
| | Arginine | R | 55 | 255 |
| | Histidine | H | 55 | 255 |

**Supplementary Table 2**. AP, F1, ACC, and AUC metrics on multi-label classification results of AMP inhibition activity against six types of drug-resistant bacteria.

| Method | AP | F1 | ACC | AUC | Total score |
|---|---|---|---|---|---|
| 3D voxel coloring+ Res-Conv Net | 0.259925139<br>*0.004475841* | **0.337807506**<br>*0.006178744* | 0.621577001<br>*0.006897216* | 0.590515614<br>*0.008653167* | **394.04** |
| 3D voxel coloring+ SwinUNETR Net | 0.211246929<br>*0.010777744* | 0.251092988<br>*0.004265785* | 0.629766214<br>*0.002313509* | 0.51133182<br>*0.013491204* | 338.32 |
| GAT | 0.250511837<br>*0.004487868* | 0.291397423<br>*0.004208417* | 0.630599976<br>*0.003318742* | 0.578253043<br>*0.008026728* | 376.18 |
| GCN | 0.269195271<br>*0.006371139* | 0.306131417<br>*0.004761229* | 0.628041649<br>*0.00592154* | 0.596994674<br>*0.00631948* | 390.22 |
| Graphsage | 0.258064944<br>*0.005238844* | 0.305261093<br>*0.002117538* | 0.627641571<br>*0.00644122* | 0.588542104<br>*0.008233152* | 384.35 |

**Explanatory note:** The values in non italics and italics represent the arithmetic mean and standard deviation calculated by 5-fold cross validation, respectively. Bold and underline indicate significant superiority over other methods. The total score is the sum of the normalized scores for each metric.

**Supplementary Table 3**. AP, F1, ACC, and AUC metrics on four antimicrobial mechanisms of AMPs.

| Method | AP | F1 | ACC | AUC | Total score |
|---|---|---|---|---|---|
| 3D voxel coloring+ Res-Conv Net | **0.399924028** | **0.422019643** | **0.910530376** | 0.653753293 | **389.98** |
| | *0.018169467* | *0.010518749* | *0.023418329* | *0.008247621* | |
| 3D voxel coloring+ SwinUNETR Net | 0.23511011 | 0.230502197 | 0.547574234 | 0.532412255 | 246.82 |
| | *0.002949666* | *0.000419831* | *0.001460559* | *0.022742496* | |
| GAT | 0.228050494 | 0.20367589 | 0.589477706 | 0.708499992 | 267.54 |
| | *0.016554248* | *0.007831837* | *0.014238752* | *0.010736589* | |
| GCN | 0.231187326 | 0.200994244 | 0.570695889 | **0.726564765** | 268.11 |
| | *0.026220621* | *0.019589181* | *0.023942926* | *0.013416302* | |
| Graphsage | 0.244186965 | 0.214947778 | 0.577783036 | 0.717550027 | 274.21 |
| | *0.029047046* | *0.014891621* | *0.01613992* | *0.004521761* | |

**Explanatory note:** The values in non italics and italics represent the arithmetic mean and standard deviation calculated by 5-fold cross validation, respectively. Bold and underline indicate significant superiority over other methods. The total score is the sum of the normalized scores for each metric.

**Supplementary Table 4**. AP, F1, ACC, and AUC metrics on multi-abel classification of six organ toxicity of AMPs.

| Method | AP | F1 | ACC | AUC | Total score |
|---|---|---|---|---|---|
| 3D voxel coloring+ Res-Conv Net | 0.120963833<br>*0.043153423* | **<u>0.146754533</u>**<br>*0.018561946* | 0.606880653<br>*0.006405425* | **<u>0.72766937</u>**<br>*0.031139092* | **358.39** |
| 3D voxel coloring+ SwinUNETR Net | 0.046975157<br>*0.009889939* | 0.05646052<br>*0.002534988* | **<u>0.693833899</u>**<br>*0.001846985* | 0.550953752<br>*0.079354125* | 241.73 |
| GAT | 0.126055226<br>*0.009502029* | 0.079594778<br>*0.006929199* | 0.571293724<br>*0.019349975* | 0.672113431<br>*0.060373561* | 302.85 |
| GCN | 0.170549801<br>*0.023893278* | 0.091171226<br>*0.009747237* | 0.545465016<br>*0.038967986* | 0.705736637<br>*0.07824469* | 337.73 |
| Graphsage | 0.124410221<br>*0.018593974* | 0.079102798<br>*0.00657142* | 0.565607727<br>*0.024043878* | 0.683791542<br>*0.067812362* | 302.34 |

**Explanatory note:** The values in non italics and italics represent the arithmetic mean and standard deviation calculated by 5-fold cross validation, respectively. Bold and underline indicate significant superiority over other methods. The total score is the sum of the normalized scores for each metric.

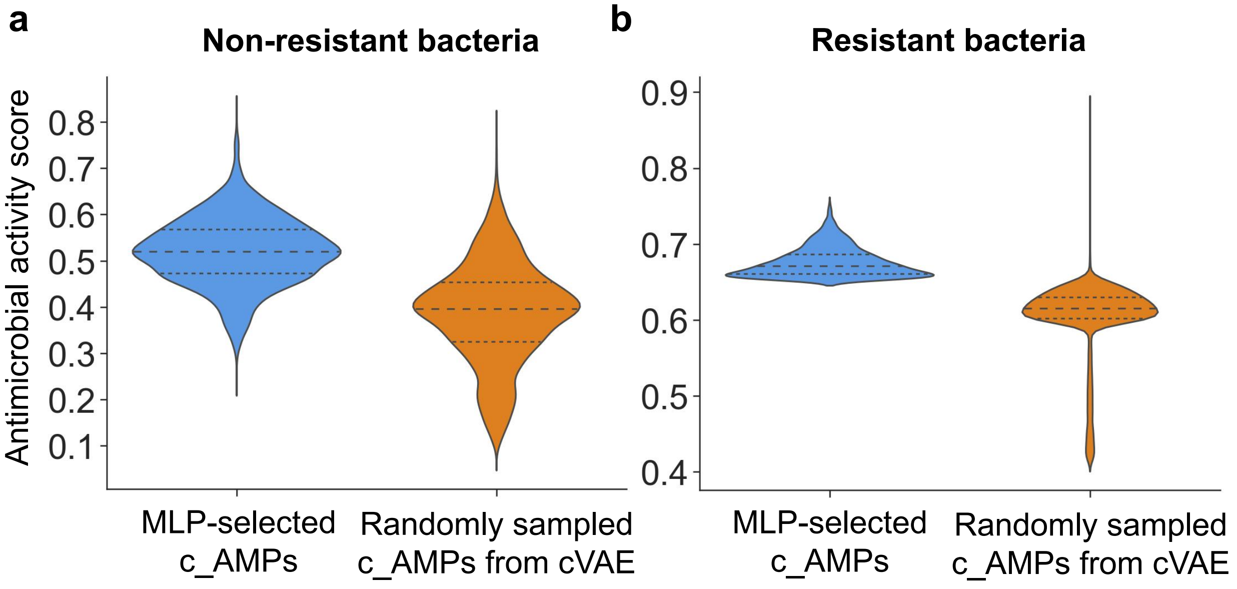

**Supplementary Fig. 2 | The predicted antimicrobial activity of the MLP-selected c_AMPs and randomly sampled peptides from cVAE.** The violin plot compares the antimicrobial activity scores of AMPs against non-resistant (a) and resistant (b) bacteria before and after screening using the MLP regression model.

**Supplementary Table 5**. The performance of multimodality is compared with using only sequence or 3D structural modalities on 3 downstream tasks.

| Task | Method | AP | F1 | ACC | AUC |
|---|---|---|---|---|---|
| Antimicrobial activity | Sequence | 0.374737072 | 0.431532663 | 0.644121861 | 0.690797079 |
| | | *0.012547968* | *0.012807786* | *0.023511166* | *0.009786222* |
| | 3D structure | 0.259925139 | 0.337807506 | 0.621577001 | 0.590515614 |
| | | *0.004475841* | *0.006178744* | *0.006897216* | *0.008653167* |
| | Multimodality | **0.396321303** | **0.453191185** | **0.680862069** | **0.705128098** |
| | | *0.004958185* | *0.005032161* | *0.007198217* | *0.006313527* |
| Mechanism | Sequence | 0.456729341 | 0.471695554 | 0.812078977 | 0.687724769 |
| | | *0.016809067* | *0.010175339* | *0.019777163* | *0.024421745* |
| | 3D structure | 0.399924028 | 0.422019643 | 0.910530376 | 0.653753293 |
| | | *0.018169467* | *0.010518749* | *0.023418329* | *0.008247621* |
| | Multimodality | **0.479836106** | **0.53599925** | **0.924816084** | **0.709135163** |
| | | *0.022431913* | *0.013762823* | *0.007993991* | *0.020707727* |
| Toxic | Sequence | **0.21297538** | 0.172096831 | 0.660560822 | 0.768246722 |
| | | *0.033818276* | *0.02836408* | *0.009581227* | *0.024093686* |
| | 3D structure | 0.120963833 | 0.146754533 | 0.606880653 | 0.72766937 |
| | | *0.043153423* | *0.018561946* | *0.006405425* | *0.031139092* |
| | Multimodality | 0.208644283 | **0.182192177** | **0.677439606** | **0.772906351** |
| | | *0.057343069* | *0.036392401* | *0.007806792* | *0.019372936* |

**Explanatory note:** The values in non italics and italics represent the arithmetic mean and standard deviation calculated by 5-fold cross validation, respectively. Bold indicate superiority over other methods.

**Supplementary Table 6**. Experimental MICs of randomly sampled 10 peptides from cVAE and the top-10 c_AMP selected from the regression moudle (3 biologically independent replicates).

| Method | Sequence | MIC (μg/ml) | | |
|---|---|---|---|---|
| | | ***S. aureus*** **CMCC26003** | ***E.coli*** **CICC21530** | ***A. baumannii*** **ATCC19606** |
| Randomly sampled 10 peptides from cVAE (Generation) | G_1 | >256 | >256 | >256 |
| | G_2 | >256 | >256 | >256 |
| | G_3 | >256 | >256 | >256 |
| | G_4 | >256 | >256 | >256 |
| | G_5 | >256 | >256 | >256 |
| | G_6 | >256 | >256 | >256 |
| | G_7 | >256 | >256 | >256 |
| | G_8 | >256 | >256 | >256 |
| | G_9 | >256 | >256 | >256 |
| | G_10 | >256 | >256 | >256 |
| Regression moudle-selected 10 c_AMPs (Generation+ Regression) | G+R_1 | 64 | >256 | 64 |
| | G+R_2 | >256 | >256 | >256 |
| | G+R_3 | >256 | >256 | >256 |
| | G+R_4 | 128 | 128 | 64 |
| | G+R_5 | 32 | 128 | 64 |
| | G+R_6 | >256 | >256 | >256 |
| | G+R_7 | 64 | >256 | 64 |
| | G+R_8 | 64 | >256 | >256 |
| | G+R_9 | 64 | >256 | >256 |
| | G+R_10 | 8 | 16 | 16 |

**Supplementary Table 7**. Physicochemical properties of training AMPs and newly discovered AMPs using the M3-CAD pipeline.

| | | | | | |
|---|---|---|---|---|---|
| Training AMPs in QLAPD (n=9,392) | **Hydrophobic** | **Hydrophobic moment** | **Aliphatic index** | **Aromaticity** | **Alpha helix** |
| | -0.251 | -0.244 | 0.979 | 0.114 | 0.361 |
| | **Beta helix** | **Turn helix** | **Charge** | **Charge density** | **Instability index** |
| | 0.174 | 0.220 | 3.952 | 0.312 | 36.825 |
| M3-CAD AMPs (n=1,000) | **Hydrophobic** | **Hydrophobic moment** | **Aliphatic index** | **Aromaticity** | **Alpha helix** |
| | -0.024 | -0.274 | 0.535 | 0.278 | 0.417 |
| | **Beta helix** | **Turn helix** | **Charge** | **Charge density** | **Instability index** |
| | 0.191 | 0.172 | -0.398 | 0.142 | 53.225 |

**Supplementary Table 8**. Physicochemical properties of antimicrobial peptide QL-AMP-1 and similarity to training AMPs.

| **Sequence** | **Hydrophobic** | **Hydrophobic moment** | **Aliphatic index** | **Aromaticity** | **Alpha helix** |
|---|---|---|---|---|---|
| QL-AMP-1 | -0.157 | -0.557 | 0.557 | 0.286 | 0.429 |
| **Sequence similarity** | **Beta helix** | **Turn helix** | **Charge** | **Charge density** | **Instability index** |
| < 30% | 0.214 | 0.214 | 2.76 | 0.143 | 26.486 |

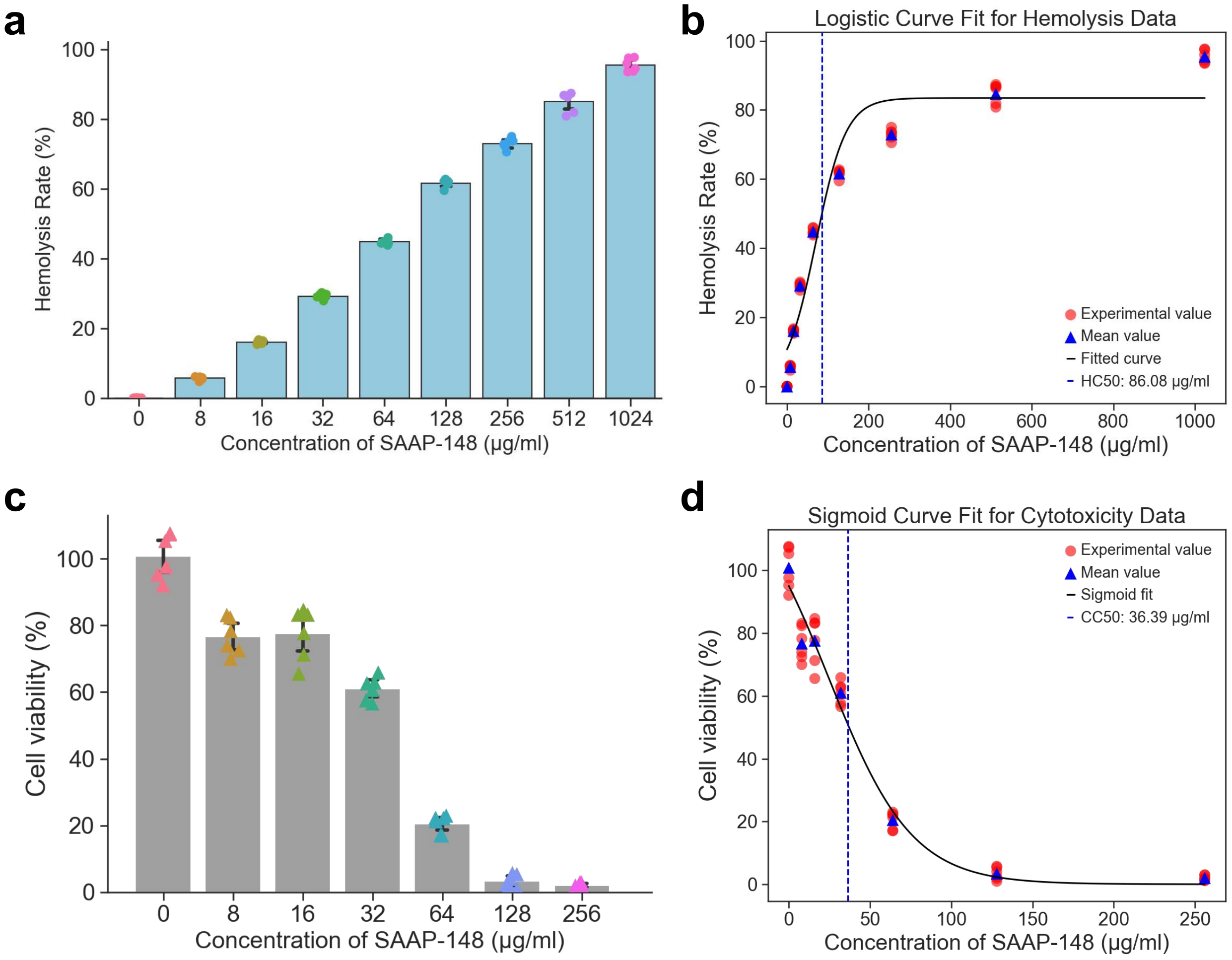

**Supplementary Fig. 3 | Experimental off-target toxicity of antimicrobial peptide SAAP-148. a-d,** Hemolysis and cytotoxicity of SAAP-148 against human red blood cells (a,b) and 293T cells (c,d) at different concentrations. HC50 (b) and CC50 (d) values are calculated by the Sigmoid function. n = 6 biologically independent replicates.

**Supplementary Table 9**. Antimicrobial activities and safe therapeutic window of QL-AMP-1 and SAAP-148.

| Species | Strain | QL-AMP-1 | | | SAAP-148 | | |
|---|---|---|---|---|---|---|---|
| | | MIC (µg / ml) | HC50 / MIC | CC50 / MIC | MIC (µg / ml) | HC50 / MIC | CC50 / MIC |
| ***S. aureus*** | ATCC25923 | 64 | 12.00 | 3.66 | 32 | 2.69 | 1.14 |
| | 181 | 128 | 6.00 | 1.83 | 32 | 2.69 | 1.14 |
| | ATCC33592 | 64 | 12.00 | 3.66 | 32 | 2.69 | 1.14 |
| | CICC25138 | 64 | 12.00 | 3.66 | 32 | 2.69 | 1.14 |
| | 129 | 64 | 12.00 | 3.66 | 32 | 2.69 | 1.14 |
| | CMCC26003 | 8 | 95.97 | 29.35 | 32 | 2.69 | 1.14 |
| ***E. faecalis*** | 152 | 128 | 6.00 | 1.83 | 32 | 2.69 | 1.14 |
| | CICC24243 | 128 | 6.00 | 1.83 | 64 | 1.345 | 0.57 |
| | ATCC51575 | 128 | 6.00 | 1.83 | 32 | 2.69 | 1.14 |
| ***E.coli*** | CICC21530 | 16 | 47.98 | 14.63 | 32 | 2.69 | 1.14 |
| | CMCC44102 | 16 | 47.98 | 14.63 | 32 | 2.69 | 1.14 |
| | 103 | 8 | 95.97 | 29.25 | 64 | 1.345 | 0.57 |
| | 110 | 16 | 47.98 | 14.63 | 32 | 2.69 | 1.14 |
| | 162 | 16 | 47.98 | 14.63 | 32 | 2.69 | 1.14 |
| | 166 | 16 | 47.98 | 14.63 | 32 | 2.69 | 1.14 |
| ***A. baumannii*** | ATCC19606 | 16 | 47.98 | 14.63 | 32 | 2.69 | 1.14 |
| | 102 | 16 | 47.98 | 14.63 | 32 | 2.69 | 1.14 |
| | 106 | 16 | 47.98 | 14.63 | 32 | 2.69 | 1.14 |
| | 114 | 32 | 23.99 | 7.31 | 32 | 2.69 | 1.14 |
| ***K. pneumoniae*** | 826 | 16 | 47.98 | 14.63 | 64 | 1.345 | 0.57 |
| | ATCC BAA-1705 | 32 | 23.99 | 7.31 | 32 | 2.69 | 1.14 |
| ***P. aeruginosa*** | 116 | 32 | 23.99 | 7.31 | 32 | 2.69 | 1.14 |
| | PAO1 | 128 | 6.00 | 1.83 | 32 | 2.69 | 1.14 |
| | 119 | 128 | 6.00 | 1.83 | 32 | 2.69 | 1.14 |
| ***Salmonella enterica*** | ATCC14028 | 32 | 23.99 | 7.31 | 64 | 1.345 | 0.57 |
| | CMCC50071 | 64 | 12.00 | 3.66 | 32 | 2.69 | 1.14 |

**Explanatory note:** HC50 and CC50 respectively refer to the half hemolytic toxicity concentration of the drug to human red blood cells and the half cytotoxicity concentration to the human embryonic kidney cell line 293T.

**Supplementary Table 10**. The secondary structure of the antimicrobial peptide QL-AMP-1 in PBS and 50% TFE solutions measured by circular dichroism spectroscopy.

| Solution | Wavelength (nm) | Helix (%) | Antiparallel (%) | Parallel (%) | Beta-Turn (%) | Rndm. Coil (%) | Total Sum (%) |
|---|---|---|---|---|---|---|---|
| PBS | 190-260 | 6.4 | 53.0 | 3.9 | 17.0 | 27.1 | 107.4 |
| | 195-260 | 7.2 | 42.5 | 5.2 | 19.1 | 32.3 | 106.3 |
| | 200-260 | 6.1 | 42.5 | 5.5 | 20.3 | 36.9 | 111.4 |
| | 205-260 | 3.4 | 44.6 | 5.2 | 19.9 | 35.1 | 108.2 |
| | 210-260 | 3.7 | 46.5 | 5.3 | 19.4 | 35.5 | 110.4 |
| 50% TFE | 190-260 | 8.1 | 46.5 | 4.0 | 18.9 | 28.2 | 105.8 |
| | 195-260 | 8.6 | 36.1 | 5.1 | 20.3 | 33.4 | 103.4 |
| | 200-260 | 7.6 | 36.1 | 5.3 | 21.4 | 36.9 | 107.3 |
| | 205-260 | 5.2 | 37.4 | 5.3 | 20.6 | 35.1 | 103.5 |
| | 210-260 | 5.6 | 39.3 | 5.4 | 19.3 | 35.2 | 104.9 |

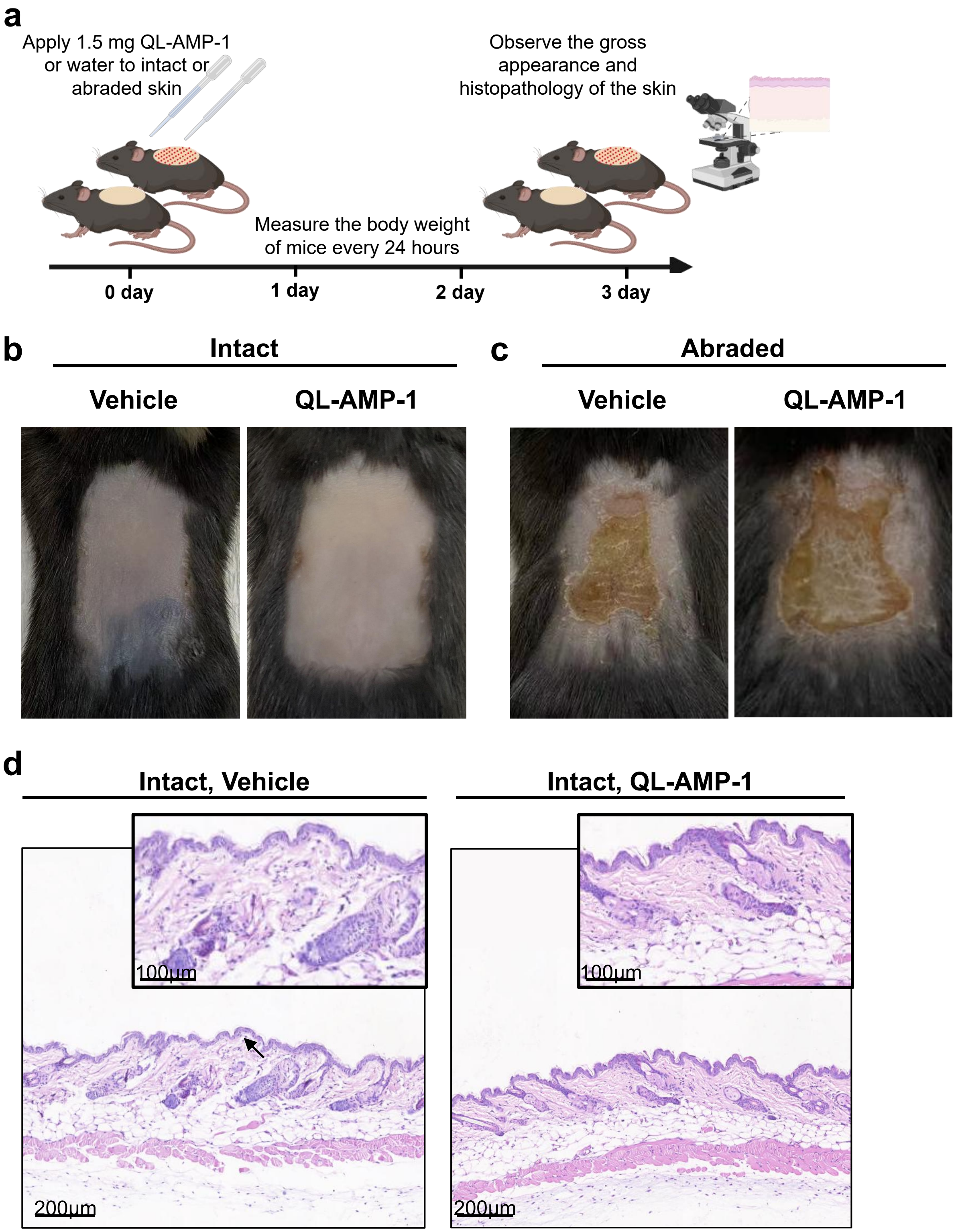

**Supplementary Fig. 4 | *In vivo* safety testing of antimicrobial peptide QL-AMP-1. a,** Schematic diagram of the animal experimental procedure for safety testing. Use sandpaper to abrade mouse skin, and then apply 5 times the therapeutic dose of QL-AMP-1 (1.5 mg) or water to the intact or abraded skin. The weight changes of the mice were measured every 24 hours. After 72 hours, the gross views of the skin were observed, and tissue biopsies were taken to observe the pathology under a microscope. **b, c,** Representative gross views of intact (b) and abraded (c) skin treated with vehicle or QL-AMP-1. **d,** Representative microscopic imaging of tissue biopsies of intact skin treated with vehicle or QL-AMP-1. n = 3 biologically independent replicates.
